# Coarse Woody Debris of Mixed Tree Species Overrides Soil Matrix Constraints on Microbial Richness and Abundance in an Ancient Broad-Leaved Forest

**DOI:** 10.64898/2026.10.09.757649

**Authors:** L.G. Khanina, M.V. Bobrovsky, D.S. Savelyev, M.N. Tutukina, T.A. Bessonova, A.G. Darchieva, P.V. Shelyakin

## Abstract

**Annotation:** The mechanisms by which coarse woody debris (CWD) affects soil microbial diversity across different spatial scales and forest disturbance regimes remain poorly understood. This study investigated the impact of CWD from multiple tree species on forest floor bacterial and fungal communities under varying landscape-scale conditions, specifically comparing old-growth forests situated on Phaozems and Retisols with mass-windthrow areas located on loamy Retisols and sandy Podzols. We analyzed both chemical compound and metagenomic profiles of 63 CWD-associated soil samples collected from the Kaluzhskie Zaseki Nature Reserve (European part of Russia). High-throughput Illumina sequencing of the 16S rRNA gene and the ITS region was used to profile both bacterial and fungal communities, respectively. Scale-dependent diversity patterns and taxonomic accumulation were assessed using GLM, NMDS, dbRDA, and iNEXT frameworks. Our findings revealed a profound spatial decoupling in forest floor microbial communities, where a localized reduction in alpha-diversity under CWD drove a sharp increase in landscape-scale gamma-diversity. While standard soil physicochemical properties rigidly dictated baseline matrix richness, CWD operated as a powerful micro-environmental hotspot overriding these constraints. Deadwood inputs filtered out dominant matrix-associated taxa, creating highly specialized micro-refugia characterized by high compositional dispersion. This niche selection was particularly pronounced beneath *Quercus* logs on Phaozems and Retisols. Furthermore, post-windthrow microbial responses were heavily filtered by wood traits and baseline soil textures. On loamy Retisols within the windthrow mosaic, this breakaway dynamic was highly taxon-specific: while *Acer* and *Fraxinus* CWD triggered partitioned, site-specific hotspot dynamics yielding an exceptionally large pool of unique ASVs for bacteria, the distinct enrichment for fungal communities was primarily driven by *Ulmus* CWD and log-free windthrow areas. Ultimately, iNEXT accumulation profiles confirmed that deadwood does not merely multiply common soil taxa, but enriches the forest landscape by injecting critical beta-level heterogeneity into the soil matrix. These insights emphasize the role of diverse CWD retention in buffering forest microbiomes against large-scale environmental disturbances across heterogeneous soil landscapes.

**Highlights:**

1. Sandy soils suppress bacterial abundance but trigger massive fungal proliferation.
2. Bacterial richness is determined by soil pH; fungal diversity responds to nitrogen.
3. Soil pH and nitrogen shift fungal composition along decoupled orthogonal axes.
4. Coarse woody debris acts as hotspots of micro-environmental diversity.
5. Post-windthrow microbial hotspots are strongly shaped by tree species.

## Introduction

Soil microbial communities play a central role in terrestrial biogeochemical cycles by driving organic matter decomposition, nutrient turnover, and carbon sequestration (Bardgett, van der Putten, 2014; Baldrian et al., 2023). Forest soils represent one of the most diverse microbial habitats on Earth with bacteria and fungi constituting the dominant components of the soil microbiome (Uroz et al., 2013; Lladó et al., 2017). Metabarcoding sequencing have greatly improved our understanding of the taxonomic composition and diversity of forest soil microbiomes, provide new opportunities to investigate the functioning and ecological stability of forest ecosystems (Uroz et al., 2013; Lladó et al., 2017; Jiao et al., 2021; Baldrian et al., 2023; Pavlopoulos et al., 2023). Nevertheless, old-growth forests remain poorly represented in microbiome studies because of their rarity and limited accessibility, despite being among the least disturbed terrestrial ecosystems and serving as important reference systems for sustainable forest management and biodiversity conservation (Lindenmayer, Franklin, 2002; Lukina et al., 2023).

Coarse woody debris (CWD, deadwood) is a key structural component of old-growth forests and performs essential ecological functions, including carbon storage, nutrient retention, and the maintenance of forest biodiversity (Stokland et al., 2012). As deadwood decomposes, it provides a substantial source of carbon and creates microsites that influence the composition and functioning of adjacent soil microbial communities. Although microbial succession within decaying wood has been extensively investigated (Bani et al., 2018; Johnston et al., 2016; Kielak et al., 2019; Qi et al., 2023), considerably less is known about how CWD affects the diversity and abundance of soil bacteria and fungi beneath fallen logs. Existing studies have examined the influence of deadwood on soil microbial communities only to a limited extent (Minnich et al., 2021; Wang et al., 2023; Błońska et al., 2024; Wang et al., 2024; Mäkipää et al., 2017) and most have focused either on a single tree species (Błońska et al., 2024; Mäkipää et al., 2017) or have not considered tree species identity (Wang et al., 2023, 2024). To our knowledge, no study has compared bacterial and fungal communities in soil beneath CWD of different tree species within multispecies old-growth broad-leaved forests.

The present study addresses this knowledge gap by investigating the abundance and diversity of soil bacterial and fungal communities in biotopes formed during autogenic succession following both mass and single windthrow events in multispecies broad-leaved forests growing on sandy and loamy soils in the Kaluzhskie Zaseki State Nature Reserve (European part of Russia). Bacterial and fungal abundance was quantified by qPCR targeting the 16S rRNA V4 region and ITS 3-4, and community diversity was assessed using metabarcoding. We tested four general hypotheses: (1) soil texture and localized micro-habitats (CWD) exert distinct, kingdom-specific constraints on absolute microbial abundance; (2) local microbial richness (alpha diversity) is filtered by different macro-chemical drivers for bacterial and fungal kingdoms; (3) soil physical and chemical properties constrain bacterial and fungal community structures (beta diversity) along independent, multi-dimensional coordinates; and (4) the micro-spatial heterogeneity sustained by mixed-species CWD acts as an ecosystem-level override, amplifying cumulative regional pools (gamma diversity) despite baseline soil constraints. This study extends our previous analysis of bacterial communities in the same forest biotopes (Savelyev et al., 2026) by incorporating fungal communities and examining the responses of soil microbiomes to contrasting windthrow biotopes, CWD of different tree species, and soil properties.

## 2. Methods

### 2.1. Study area

The study was conducted in the southern section of the Kaluzhskie Zaseki State Nature Reserve (53°30′-53°39′ N, 35°37′-35°49′ E), located in the northwestern part of the Central Russian Upland within the upper Oka River basin (Popadiyuk et al., 1999; Smirnova et al., 2017). The Reserve encompasses extensive broad-leaved forests that have existed since the Trans-Oka Abatis Line of the Muscovite State (16th-17th centuries) and are now among the best-preserved old-growth forests in the temperate zone of European Russia (Smirnova et al., 2017). The climate of the region is moderately continental. According to data from the nearest meteorological station, Sukhinichi (65 km from the study area), the mean annual air temperature was 5.9 °C in 1991-2020, increasing to 6.3 °C in 2006-2020 (Bulygina et al., 2022). Mean monthly temperatures range from -7.3 °C in January to 18.8 °C in July. The frost-free period lasts approximately 140 days, and the growing season (mean daily temperature > 5 °C) about 180 days. Mean annual precipitation, with all corrections, is 662 mm (Ilyin et al., 2022).

The landscape is a gently sloping upland with an erosional surface, located mainly at elevations of 150-250 m a.s.l. Pre-Quaternary deposits are Early Cretaceous clays, sands and sandstones. Quaternary deposits are 10-20 m thick and are dominated by Early-Middle Pleistocene till of the Don glacial complex, Middle Pleistocene fluvioglacial deposits and Late Pleistocene loess-paleosol strata (Novenko et al., 2019). Loess-like loamy deposits are dominated by Albic Retisols and Albic Luvisols, with Luvic Phaeozems occurring locally. Sandy deposits are occupied by Albic and Entic Podzols (IUSS, 2022; Bobrovsky, 2010). Forests on loamy deposits belong to the association *Aceri campestris-Tilietum cordatae* ass. nov. hoc loco (Zaugolnova, Braslavskaya, 2003), characterized by high constancy of *Acer campestre*, *Euonymus europaeus* and *Allium ursinum*. Broad-leaved forests on sandy soils belong to the association *Querco-Tilietum cordatae* (Laivinsh 1986 ex Laivinsh in Solomeshch et al., 1993), characterized by high constancy of nemoral species and rare occurrence of boreal species (*Gymnocarpium dryopteris*, *Luzula pilosa*, *Maianthemum bifolium*, *Phegopteris connectilis*, etc.). According to the EUNIS habitat classification (Chytrý et al., 2020), both forest types belong to the *Quercus* mesic deciduous forest habitat (T1E). The overstorey is formed, in different proportions at different sites, by *Quercus robur*, *Fraxinus excelsior*, *Tilia cordata*, *Ulmus glabra*, *Acer platanoides*, *A. campestre*, *Populus tremula*, *Betula pubescens* and *B. pendula*; *Picea abies* and *Pinus sylvestris* also occur in the canopy. Nemoral species such as *Aegopodium podagraria*, *Pulmonaria obscura*, *Galeobdolon luteum*, *Glechoma hirsuta* and *Dryopteris carthusiana* are common in the ground vegetation.

The study aimed to explore the soil of multispecies broad-leaved forests affected by large-scale or single-tree windthrow. We selected sites differing in soil texture, including both an old-growth forest with single-tree gap dynamics (Shashkov et al., 2022) and mature forests that experienced mass windthrow in the summer of 2006 (Bobrovsky, Stamenov, 2020). Soil was sampled in four biotopes: two in the mass windthrow area, on sandy Podzols (PZ) and loamy Retisols (Ret_N), and two in the old-growth forest with isolated windthrows, on loamy Retisols (Ret_T) and Phaeozems (PH) (Figs. 1, 2). In PZ, the sandy (> 0.05 mm) and silty (0.001-0.05 mm) fractions accounted for 89% and 10% of particles, respectively, and the clay fraction (< 0.001 mm) for only 1%. In the loamy biotopes, the mean sand:silt:clay ratio was 24:69:7, although the fraction ratios differed significantly among biotopes (Table S1; Khanina, Bobrovsky, 2024).

**Fig. 1.**
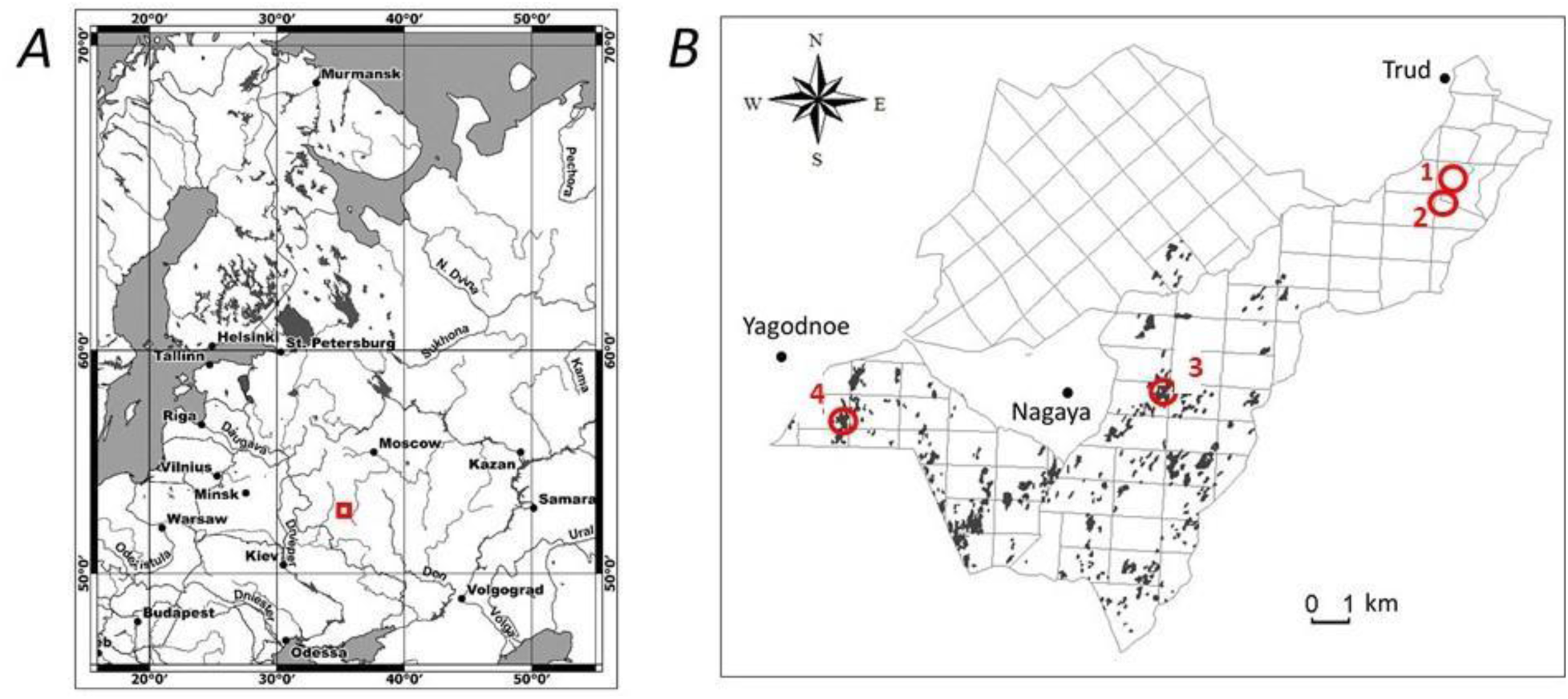
Study area (*A*, red square) and biotopes (*B*, red circles) in the Southern section of the Kaluzhskie Zaseki Reserve: 1 and 2 old-growth forest (over 300 years old) with isolated windthrows on Phaozems (PH) and Retisols (Ret_T); 3 and 4 mass windthrow of 2006 in broad-leaved forest on Retisols (Ret_N) and Podzols (PZ). Black spots are patches of the 2006 mass windthrow. From (Bobrovsky, Stamenov, 2020) with modifications.

**Fig. 2.**
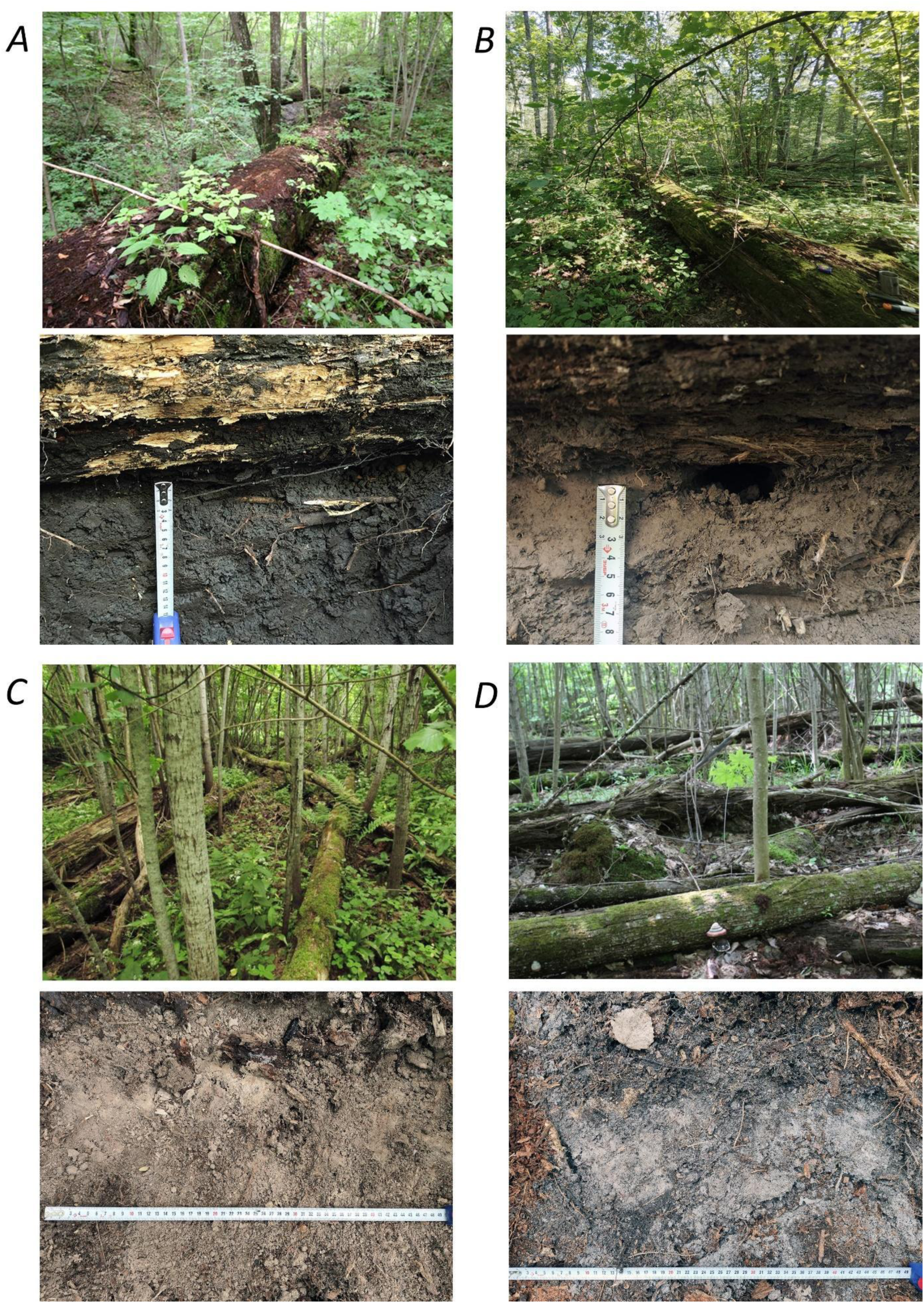
Images of the study biotopes and soil sampling sites. (*A* and *B*) Large fallen oaks in the old-growth broad-leaved forest on Phaozem (PH) and Retisol (Ret_T); (*C* and *D*) mass windthrow areas on Retisol (Ret_N) and Podzol (PZ).

### 2.2. Collection of soil samples

Soil samples for metagenomic DNA isolation were collected in 2024 from the top 5 cm of the A horizon (excluding the litter layer) beneath coarse woody debris (CWD) of seven tree species: *Acer platanoides*, *Fraxinus excelsior*, *Populus tremula*, *Tilia cordata*, *Ulmus glabra*, *Quercus robur* and *Picea abies*. All selected logs were at decomposition stage 3 (on a five-point scale) and lay in close contact with the soil surface. Samples were also collected at two sites outside CWD: a mass windthrow area (“Windthr”) and a background forest (“Forest”). At least three replicates were sampled per site, taking into account the tree species of the fallen log. In total, 63 samples were collected from nine microsites (seven CWD species and two sites outside CWD) across four biotopes (Table 1). Samples were collected with sterilized instruments, placed in sterile Whirl-Pak bags, frozen at -20 °C, and transported to the laboratory in a cooler without thawing. Soil samples for chemical analysis were collected simultaneously at the same sites.

**Table 1.**
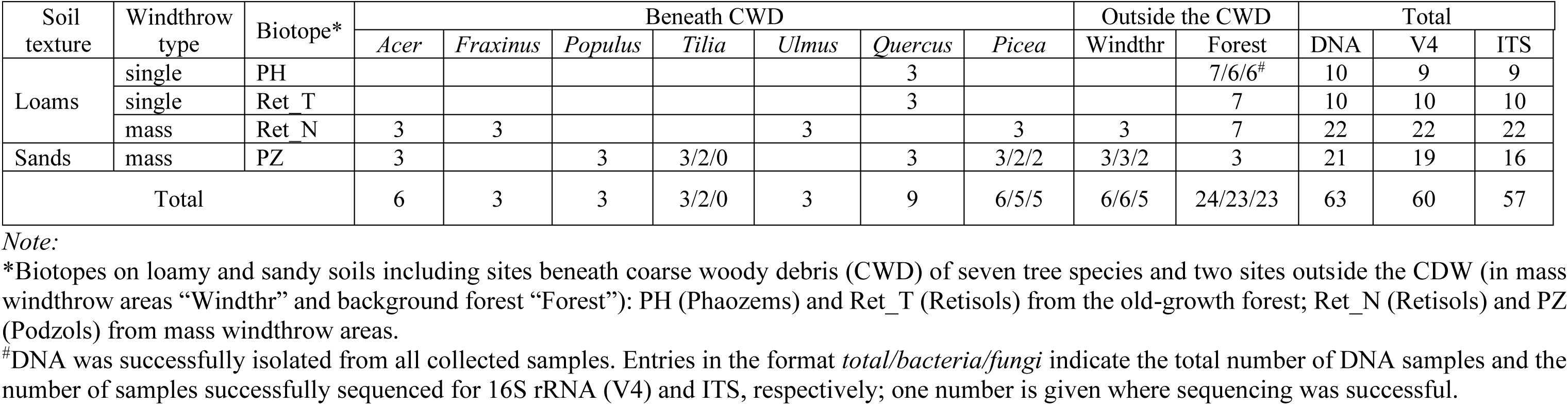
Number of soil samples collected for DNA isolation and successfully sequenced for the bacterial 16S rRNA (V4) and fungal ITS genes across four forest biotopes and nine sites (from Savelyev et al., 2026 with modifications)

**Table 2.** Soil chemical characteristics across the studied biotopes

| Biotope | n | C (%) | N (%) | C/N | pH (H<sub>2</sub>O) | CEC (cmol<sub>c</sub>/kg) | P<sub>2</sub>O<sub>5</sub> (mg/kg) | K<sub>2</sub>O (mg/kg) |
| --- | --- | --- | --- | --- | --- | --- | --- | --- |
| PH | 10 | 5.23 ± 0.27<sup>a</sup> | 0.44 ± 0.02<sup>a</sup> | 11.88 ± 0.14<sup>a</sup> | 5.88 ± 0.10<sup>ab</sup> | 33.5 ± 2.2<sup>a</sup> | 53.6 [49.4; 53.6]<sup>a</sup> | 200.0 [200; 238.8]<sup>a</sup> |
| Ret_T | 10 | 3.07 ± 0.20<sup>b</sup> | 0.24 ± 0.02<sup>b</sup> | 12.70 ± 0.34<sup>a</sup> | 5.42 ± 0.09<sup>bc</sup> | 15.8 ± 2.2<sup>c</sup> | 33.0 [28.1; 33.0]<sup>bc</sup> | 95.0 [95; 121.3]<sup>b</sup> |
| Ret_N | 22 | 2.75 ± 0.11<sup>b</sup> | 0.23 ± 0.01<sup>b</sup> | 11.99 ± 0.20<sup>a</sup> | 6.30 ± 0.14<sup>a</sup> | 23.6 ± 1.5<sup>b</sup> | 40.6 [28.7; 56.3]<sup>ac</sup> | 165.0 [165; 327.5]<sup>a</sup> |
| PZ | 21 | 3.37 ± 0.23<sup>b</sup> | 0.26 ± 0.02<sup>b</sup> | 13.07 ± 0.46<sup>a</sup> | 5.05 ± 0.14<sup>c</sup> | 10.4 ± 1.5<sup>c</sup> | 18.9 [14.6; 26.5]<sup>b</sup> | 80.0 [65.0; 105.0]<sup>b</sup> |

Note:
Data are presented as mean ± standard error for C, N, C/N, pH, and CEC and as median [25th; 75th percentiles] for AP and AK. Different superscript letters within a column indicate statistically significant differences between biotopes based on parametric (Tukey-Kramer) or non-parametric (Dunn's with Holm's adjustment) post-hoc tests ( $p < 0.05$ ).

### 2.3. Chemical soil analysis

Soil samples were air-dried and sieved (1 mm mesh). Total carbon (C, %) and nitrogen (N, %) were determined in samples ground to a powder using a Vario EL Cube CHNS analyzer (Elementar Analysensysteme GmbH, Germany). Soil pH was measured potentiometrically in an aqueous suspension at a soil:distilled water ratio of 1:2.5 (Sartorius Basic Meter PB-11, Germany). Available phosphorus (P₂O₅) and potassium (K₂O) (AP and AK, respectively) were extracted with 0.2 M HCl and determined colorimetrically (UNICO-1200 spectrophotometer, USA) and by flame photometry (BWB-XP, Performance Plus, BWB Technologies, UK), respectively. Cation exchange capacity (CEC) was determined by the BaCl₂ method (Hendershot, Duquette, 1986).

### 2.4. Amplification and sequencing of the V4 region of 16S rRNA and ITS 3-4 region

Metagenomic DNA was isolated from each soil sample using DNeasy PowerSoil Pro Kit (Qiagen, Germany) according to the manufacturer’s instruction, and stored at -20°C until use. The isolated DNA was used both for quantitative real-time PCR (qPCR) to determine bacterial and fungal abundance based on threshold cycle (Ct) values, and for microbial community profiling via high-throughput sequencing.

For community analysis, the V4 variable region of the 16S rRNA gene (for bacteria) and the 3-4 subregion of the internal transcribed spacer (ITS) (for fungi) were amplified. Amplicons were generated using standard primers adapted for the Illumina platform, which contained the appropriate Illumina locus-specific overhang sequences and high-fidelity Q5 polymerase (Q5 High-Fidelity 2X Master Mix, NEB, United States). The V4 variable region of the 16S rRNA gene was amplified with the primers 515F 5ʹ-GTGCCAGCMGCCGCGGTAA-3ʹ and 806R 5ʹ-GGACTACHVGGGTWTCTAAT-3ʹ and 3-4 subregion of the ITS with the primers ITS3 GCA TCG ATG AAG AAC GCA GC and ITS4 TCC TCC GCT TAT TGA TAT GC. The PCR conditions were individually optimized for each primer set; the general program included initial denaturation at 95°C for 2 min, followed by 27 (V4) or 32 (ITS) cycles of denaturation (95°C, 25 s), primer annealing (55 (ITS) or 58°C (V4), 30 s), and elongation (72°C, 40 s), with a final elongation at 72°C for 3 min. A negative control without DNA was included in each batch to exclude possible contamination.

The obtained amplicons were purified using AMPure XP beads (1:0.9 v/v, Beckman Coulter, United States), and their concentration was measured with a Qubit fluorimeter (Thermo Fisher Scientific, United States). Index PCR was then carried out using the Nextera XT Index Kit (Illumina, United States) and Q5 High-Fidelity 2X Master Mix according to the manufacturer’s protocol. The resulting amplicon libraries were purified on AMPure XP beads (1:1.15 v/v) and quantified using Qubit. Paired-end sequencing (2 × 250 bp) of the libraries was carried out at the Genomics and Bio-imaging Core Facility (Skoltech, Russia), using a MiSeq system and a MiSeq Reagent Kit v2 (500 cycles, Illumina).

#### Quantitative PCR

Amplification was run on a DT-lite thermal cycler (DNA-Technology, Russia) using primers without Illumina adaptors and the same programs as for community profiling but without final elongation. Fluorescence signal detection was performed at the end of each cycle, followed by a melting curve analysis to confirm product specificity. A negative control without DNA was included in each batch to exclude possible contamination. Two technical replicates were used for each sample from each batch. The numbers of doubling PCR cycles required to detect bacterial (*Ct*_V4) and fungi (*Ct*_ITS) DNA were estimated; the lower the number, the higher was the DNA content.

### 2.5. Analysis of sequencing data

Quality of the sequences obtained was assessed using FastQC v. 0.12.0. Sequencing data were analyzed using QIIME2 v. 2025.9 (Bolyen et al., 2019). DADA2 v. 2025.10.0.dev0 was used to correct sequencing errors, to merge paired reads, to eliminate chimeric sequences, and to build an amplicon sequence variant (ASV) tables (Callahan et al., 2016).

For analysing 16S rRNA standard parameters of the DADA2 function were changed to obtain a greater number of reads at the taxonomic assignment step. We used qiime dada2 denoise-paired with the following modified parameters: truncLen_left = 190, truncLen_right = 120, maxEE_left = 3, maxEE_right = 3, trim_left = 30, min_overlap = 8, and max_mismatch = 1; other parameters were default. The mean read number per sample was approximately 35 000 (minimum, 8000; maximum, 105 000) before data filtering. Fewer than ten reads were obtained for three samples; the respective samples were excluded from further analysis. On average, 43% of all reads passed through all filtration steps in DADA2; after data filtering the mean read number was approximately 15 000 in each sample. Taxonomic assignments of reads were established and included in the ASV table with the use of the q2-feature-classifier plugin and the classifier 2024.09.backbone.v4.nb.sklearn-1.4.2.qza, which was preliminarily trained using reference sequences from the GreenGenes2 database (McDonald et al., 2024). The classifier was available at the QIIME2 site. The final ASV table constructed for 60 samples and the taxonomic assignments and metadata of the samples were converted to a phyloseq object in R v. 4.5.1 2025-06-13 ucrt (R Core Team, 2025) for further analysis. The function rarefy_even_depth of the package Phyloseq v. 1.52.0 (McMurdie et al., 2013) was used to randomly rarefy the data to an even number of reads per sample to account for differences in sequencing depth (the final rarefaction depth was 4554); the number of reads from a single ASV was converted to their proportion per sample by using the function transform_sample_counts(phyloseq.object, function(OTU) OTU/sum(OTU)).

ITS sequences were denoised with the qiime dada2 denoise-single method (QIIME 2 v. 2025.9; Bolyen et al., 2019) using --p-trim-left 0, --p-trunc-len 268 and --p-max-ee 3; all other parameters were left at their defaults. Before filtering, the mean number of reads per sample was ≈15000 (min 2500, max 40000). Three of the 60 samples yielded fewer than ten filtered reads and were excluded, leaving 57 samples for fungal analyses (three fewer than in the 16S dataset). On average, 47% of reads passed filtering, denoising and chimera removal, leaving a mean of ≈7000 reads per sample. Taxonomy was assigned with qiime feature-classifier classify-sklearn (scikit-learn 1.4.2, confidence 0.7) using a naïve Bayes classifier trained with fit-classifier-naive-bayes on UNITE v10.0 (2025-02-19) reference sequences of species hypotheses (SHs) clustered at 99% similarity (Abarenkov et al., 2025). The ASV table, taxonomy and metadata for 57 samples were combined into a phyloseq object in R v4.5.1 (R Core Team, 2025). Data were rarefied to 2220 reads per sample with rarefy_even_depth (phyloseq v1.52.0 (McMurdie et al., 2013); replace = FALSE), and ASV counts were then converted to per-sample proportions with transform_sample_counts(phyloseq.object, function(OTU) OTU/sum(OTU)).

Relative abundances of major taxonomic groups were visualized using bar plots aggregated at the levels of class (for bacteria), order (both for bacteria and fungi), and genus (for fungi) to provide an overview of community dominance patterns across biotopes and sites analyzed. Importantly, in this article, the term ’bacteria’ is used in a broad sense; it encompasses archaeal representatives, as they share similar morphology and perform overlapping ecological functions alongside true bacteria. The finalized ASV abundance matrices (X_V4 for bacteria and X_ITS for fungi) generated using Phyloseq were used to calculate local alpha diversity indices. Fungal and bacterial alpha diversities per individual sample were quantified as the number of observed ASVs, Chao1, Shannon, and inverse Simpson diversity indexes. All alpha diversity metrics and taxonomic matrices were subsequently exported for formal downstream statistical modeling.

### 2.6. Statistical Modeling of Microbial Abundance and Diversity

All statistical analyses were performed in the R statistical environment (v4.4.0) (R Core Team, 2025). To check the hypotheses, all outputs were grouped by the four biotopes and by nine sites. Significance of differences in chemical characteristics of samples and alpha-diversity metrics were evaluated by a series of one-way ANOVA with subsequent pairwise comparisons of the means (in the case of significant differences in a group) by Tukey test. The homogeneity of the variances was checked using the Levene test in the car package.

To evaluate the absolute abundance of the bacterial and fungal communities, general linear models (GLMs) were fitted using the qPCR threshold cycle (*Ct* for the 16S rRNA gene V4 region and ITS, respectively) as continuous response variables; lower *Ct* values indicated higher initial gene copy numbers. To enable a direct comparison between the two microbial groups, both communities were initially evaluated using a baseline additive model that incorporated soil texture (Loam vs. Sand) and spatial position relative to deadwood (Beneath CWD vs. Outside CWD) as categorical predictors. Subsequently, a comprehensive model selection procedure based on Akaike information criterion (AIC) and adjusted *R*^2^ was performed to optimize model fit for each community. These criteria prioritized parsimonious models incorporating categorical environmental variables over models over-saturated with soil chemical properties. For the bacterial community, the initial baseline additive model remained the most parsimonious and top-ranked fit. In contrast, this additive model performed poorly for the fungal community. Therefore, the final top-ranked model for fungi was expanded to include the interaction between the four biotopes and nine sites, which significantly improved model fit.

To evaluate the effects of soil chemical properties and type of soil texture (Loam vs. Sand) on bacterial and fungal alpha diversities, we constructed generalized linear models (GLMs) where bacterial and fungal alpha diversities were quantified as the observed number of ASVs per sample. Given that the ASV count data both for bacterial and fungi exhibited substantial overdispersion (residual deviance significantly exceeded the degrees of freedom under a standard Poisson model), Negative Binomial regression models with a log-link function (glm.nb from the MASS package) were deployed to ensure robust standard errors and eliminate Type I errors. Prior to model fitting, all continuous environmental predictors (soil C, N, pH, CEC, AP, and AK) were standardized using Z-score transformation (mean = 0, SD = 1) to ensure comparability of their regression coefficients and stabilize model convergence. Potential multicollinearity among these variables was assessed using the Variance Inflation Factor (VIF) from the car package. A severe multicollinearity issue was detected between C and N (VIF > 35), reflecting their tightly coupled dynamics across the biotopes. To stabilize model estimates, total carbon was omitted from all final equations, which reduced the VIF values for all remaining continuous predictors below a conservative threshold of 5.1, confirming the absence of significant multicollinearity. In accordance with the principle of parsimony, Type of soil texture (Loam vs. Sand) was treated as a categorical factor with ’Loam’ set as the reference baseline (Table S1). Models fit and assumptions were verified by checking the ratio of residual deviance to residual degrees of freedom (∼1.15 both for fungi and bacteria), confirming excellent fit. Regression coefficients were exponentiated and reported as Incidence Rate Ratios (IRRs), which represent the proportional change in ASV richness per SD increase in a given predictor. Model effects were visualized using the sjPlot and ggeffects packages, with marginal effects back-transformed and plotted against the unscaled, raw environmental variables.

To visualize and formally evaluate shifts in microbial community composition (beta diversity) across sites, biotopes, and environmental gradients, a combined unconstrained and constrained ordination approach was applied. First, Non-metric Multidimensional Scaling (NMDS) with the Bray-Curtis metrics was performed using the metaMDS function in the Vegan package (Oksanen et al., 2025) to visually assess the structural distribution and dissimilarity of bacterial and fungal communities. This unconstrained approach allowed for the qualitative evaluation of microbial clustering patterns across different Biotopes, Sites (beneath CWD of seven tree Species, beyond CWD in the mass Windthrows, and in background Forest outside CWD). Differences in community composition between sample groups were tested for significance by permutational multivariate ANOVA (PERMANOVA) (Anderson, 2017) (the functions betadisper and adonis2 of the Vegan package). Second, to formally test how specific soil environmental properties and categorical filters constrain microbial beta diversity, distance-based Redundancy Analysis (dbRDA) was executed using the dbrda function in vegan. The dbRDA models crossed the same bacterial (X_V4) and fungal (X_ITS) Bray-Curtis matrices with the same set of standardized soil variables and Bedrock as a categorical factor. The global significance of the constrained models, as well as the independent, unique contribution of each predictor, was rigorously verified using marginal permutation tests (anova.cca with 999 permutations) via the by = “margin” argument to control for covariates. Notably, whilst total carbon content also showed a significant individual effect, but it was largely collinear with nitrogen content; ultimately, the model incorporating nitrogen provided a better fit.

To evaluate and compare the cumulative species pools of the study area (gamma richness) and evaluate sampling saturation, sample-size-based and coverage-based rarefaction and extrapolation curves were constructed using the iNEXT package (Chao et al., 2014; Hsieh et al., 2016). The analysis was executed at two distinct hierarchical levels for both the bacterial (X_V4) and fungal (X_ITS) datasets: (1) macro-environmentally, by aggregating samples within each Biotope, and (2) macro-ecologically, by profiling individual locations (Site) nested within those biotopes to capture internal spatial heterogeneity. The site-level analysis was explicitly linked with CWD inputs, enabling the evaluation of cumulative diversity pools sustained by different tree species of decaying logs compared to control forest matrices. Microbial diversity profiles were comprehensively quantified using Hill numbers of three orders: *q* = 0 (observed ASV richness), *q* = 1 (Shannon diversity, exponential of Shannon index), and *q* = 2 (Simpson diversity, inverse of Simpson concentration) to evaluate changes across both rare and dominant taxonomic fractions. Asymptotic diversity estimates, along with their associated 95% confidence intervals, were calculated via bootstrapping (999 replications) to predict total expected richness at full sampling saturation. Non-overlapping 95% confidence intervals between the interpolated and extrapolated curves were utilized as a rigorous, conservative criterion to determine statistically significant differences in microbial cumulative reservoirs across the analyzed biotopes and individual sites.

## 3. Results

### 3.1. Soil Chemical Properties

The investigated biotopes exhibited pronounced variations in their soil chemical profiles (Table 2). The contents of total carbon (C) and total nitrogen (N) in the soil showed very similar patterns across the study sites, with significantly higher stocks observed in the PH biotope (5.23 ± 0.27% and 0.44 ± 0.02%, respectively) compared to all other variants (*p* < 0.0001). Conversely, the carbon-to-nitrogen (C/N) ratio remained statistically uniform across all four biotopes, indicating a stable baseline quality of soil organic matter in the biotopes studied.

**Table 2.** ANOVA results for the general linear models predicting bacterial and fungal qPCR *Ct* values (Type II)

| Community and Source | df | Sum Sq | Mean Sq | F-value | p-value |
| --- | --- | --- | --- | --- | --- |
| Bacterial community (*Ct_V4*) |  |  |  |  |  |
| Bedrock: Loam vs. Sand | 1 | 2.87 | 2.87 | 18.79 | < 0.001 |
| Place: Beneath CWD vs. Outside the CWD | 1 | 1.05 | 1.05 | 6.87 | 0.011 |
| Residuals | 60 | 9.15 | 0.15 |  |  |
| Fungal community (*Ct_ITS*) |  |  |  |  |  |
| Biotope: PH / PZ / Ret_N / Ret_T | 3 | 16.76 | 5.59 | 10.30 | < 0.001 |
| Site: beneath CWD of seven tree species and two sites outside the CWD | 8 | 12.34 | 1.54 | 2.84 | 0.012 |
| Biotope:Site | 5 | 7.66 | 1.54 | 2.83 | 0.026 |
| Residuals | 46 | 24.95 | 0.54 |  |  |

Soil pH and cation exchange capacity (CEC) displayed wide and consistent gradients across the biotopes. The values shifted from moderately acidic conditions with lower exchange capacities in the sandy PZ biotope (pH 5.05 ± 0.14; CEC 10.4 ± 1.5 cmol_c/kg) to slightly acidic, high-capacity systems in the loamy Ret_N (pH 6.30 ± 0.14) and PH (CEC 33.5 ± 2.2 cmol_c/kg) variants, which significantly outperformed the other two (*p* < 0.05). Non-parametric analyses for mobile nutrients demonstrated a clear difference among the sites. Mobile phosphorus (AP) and available potassium (AK) concentrations were significantly lower in the PZ biotope (medians of 18.9 mg/kg and 80.0 mg/kg, respectively) relative to PH and Ret_N (*p* < 0.05), which formed a distinct macronutrient-rich group. Taken together, these results outline PH and Ret_N as nutrient-dense and highly buffered loamy systems. Notably, although the PZ biotope (Podzol) developed on sandy substrates showed the lowest absolute chemical values, its properties — including a moderate pH of 5.05, stable C/N ratio, and substantial organic matter retention (C: 3.37%) — demonstrate a remarkably well-functioning and fertile state for a sandy soil ecosystem, contrasting with typical nutrient-depleted podzolic baselines.

### 3.2. Bacterial and Fungal DNA Loads (qPCR)

Initial baseline modeling revealed that soil texture acted as a primary driver filtering both bacterial and fungal DNA abundance, but in opposing directions (Fig. 3). In sandy matrices compared to loam, bacterial abundance was systemically suppressed (Estimate = 0.47 ± 0.11, *p* < 0.001), whereas fungal abundance was significantly enhanced (Estimate = -1.22 ± 0.23, *p* < 0.001). Furthermore, spatial position relative to CWD impacted the communities differently: *Ct_*V4 values were systematically lower outside the CWD than beneath it (Estimate = -0.27 ± 0.10, *p* = 0.011; model adjusted *R*^2^ = 0.34) indicating a suppression of bacterial abundance beneath deadwood, while total fungal abundance showed no significant overall response to binary site position (*p* = 0.07).

**Fig. 3.**
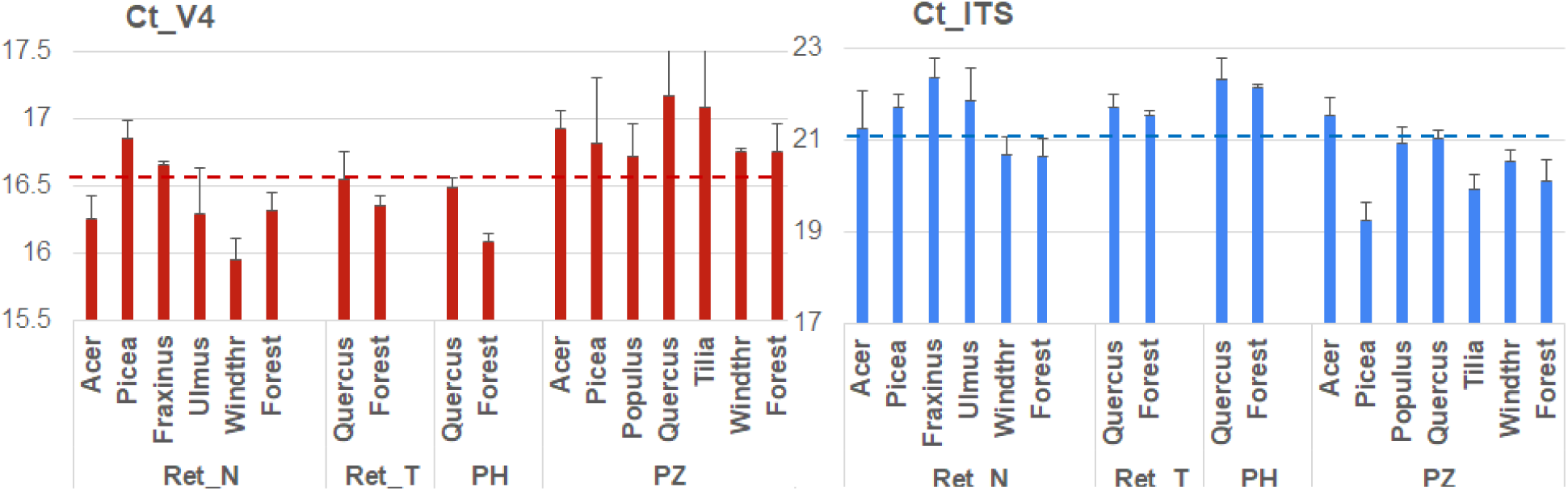
The number of doubling PCR cycles required to detect bacterial and fungal DNA signal (*Ct*_V4 and *Ct*_ITS). Lower values suggest higher target DNA contents. The horizontal lines represent the mean *Ct* values for the samples.

To better resolve the specific drivers of fungal dynamics, a comprehensive interactive model was deployed, which accounted for 50.2% of variance (*F* (16, 46) = 4.91, *p* < 0.001), adjusted *R*^2^ = 0.502). ANOVA confirmed that fungal *Ct* values were highly dependent on the synergistic interaction between Biotope and the Site (*p* = 0.026) (Table 2). Regression coefficients revealed that *Tilia* debris exerted a significant individual stimulating effect on fungal abundance across biotopes (Estimate = -1.60 ± 0.60, *p* = 0.011). Crucially, the interaction was most pronounced for *Picea* logs within the PZ biotope, demonstrating a strong synergistic effect that exceptionally amplified fungal gene copy numbers under spruce debris in sandy Podzols (Estimate = -2.77 ± 0.85, *p* = 0.002).

### 3.3. Taxonomic Composition of Bacterial and Fungal Communities

In total, 19287 bacterial and 1565 fungal ASVs were detected in the samples. ASV lists of individual samples and metadata are available from the Figshare repository (Savelyev, 2025).

*Alphaproteobacteria* (the order Rhizobiales) was the most abundant bacterial class in the majority of samples (Fig. 4A). The class *Verrucomicrobiae* (the orders *Chthoniobacterales* and *Vicinamibacteriales*) was the second most abundant. High abundance was also observed for bacteria of the classes *Planctomycetia* (the orders *Gemmatales, Isosphaerales*, and *Pirellulales*), *Gammaproteobacteria* (*Burkholderiales*), *Vicinamibacteria* (*Vicinamibacteriales*), and *Actinomycetes* (*Mycobateriales* and *Streptosporangiales*).

**Fig. 4.**
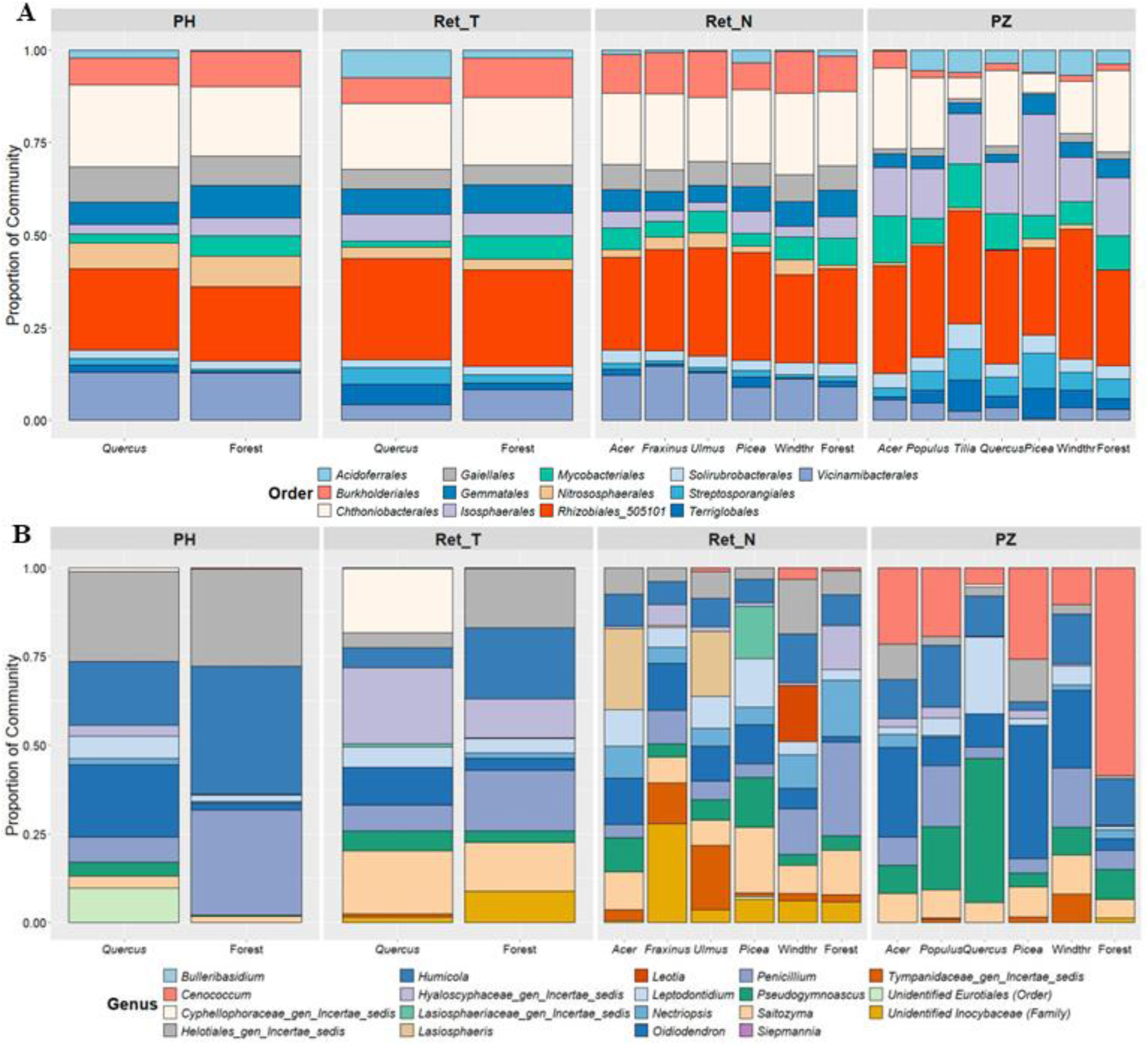
Taxonomic compositions of bacterial (A) and fungal (B) communities, averaged across all repeats at the order and genus levels, respectively. *Note:* Here and throughout the text, ’bacteria’ also refers to certain prokaryotic counterparts from the domain Archaea (e.g., *Nitrososphaerales*), due to their structural similarities and joint participation in soil processes.

The taxonomic structure of the soil bacterial communities was characterized by a highly conserved core microbiota across all studied sites and biotopes (Fig. 4A). The orders *Rhizobiales* and *Chthoniobacterales* consistently emerged as the primary co-dominants in nearly all analyzed plots. This stable taxonomic backbone was uniformly supplemented by a diverse assembly of sub-dominant groups, including *Vicinamibacteriales, Burkholderiales, Gaiellales*, and *Isosphaerales.* In the PH and Ret_N biotopes, *Burkholderiales* and *Vicinamibacteriales* maintained a steady and uniform presence across sites beneath CWD of *Acer*, *Fraxinus*, *Ulmus*, and *Picea*, as well as within the Windthrow and Forest areas where samples were taken outside the logs. Localized structural shifts were minor and restricted to specific contexts. Notably, an increased prominence of *Isosphaerales* and *Mycobacteriales* was observed in the PZ biotope (particularly under *Acer*, *Populus*, *Quercus*, and *Picea* CWD), while *Gemmatales* and the archea *Nitrososphaerales* reached higher relative abundances primarily in the PH biotope.

In contrast to the bacteria, the taxonomic composition of the fungal communities exhibited pronounced heterogeneity and distinct host-specific variations across the studied sites. The overall community structure was broadly characterized by the widespread abundance of saprotrophic microfungi *Ascomycota*, including *Penicillium (*order *Eurotiales*)*, Humicola (Sordariales), Oidiodendron (Helotiales),* and *Pseudogymnoascus (Thelebolales).* Among the dominant representatives of *Basidiomycota* are the unicellular yeasts of the genus *Saitozyma* (order *Tremellales*). The taxa are found predominantly in loamy soils. In sandy Podzols, ectomycorrhizal fungi of the genus *Cenococcum* (*Gloniales*) account for a significant proportion.

Within the PH biotope, *Helotiales* and *Oidiodendron* dominated under *Quercus* CWD, whereas *Humicola* and *Penicillium* were more prevalent in the Forest control plots outside logs. In the Ret_T biotope, sites beneath *Quercus* CWD were uniquely co-dominated by *Hyaloscyphaceae* and *Cyphellophoraceae*, while Forest sites maintained a standard profile of *Humicola* and *Penicillium*. The Ret_N biotope demonstrated the highest compositional diversity; sites beneath *Acer* and *Ulmus* CWD were heavily populated by *Lasiosphaeris* and *Leptodontidium,* while *Fraxinus* hosted unidentified *Inocybaceae*, and *Picea* exhibited an expansion of *Satiozyma* and *Lasiosphaeriaceae*. In the Windthrow area within Ret_N, *Leotia* and various *Helotiales* emerged as dominant components. The PZ biotope was heavily marked by the prevalence of the *Cenococcum* across *Acer*, *Populus*, *Picea*, and Forest sites, while *Pseudogymnoascus* and *Leptodontidium* became more prominent specifically under *Quercus* CWD here.

### 3.4. Patterns and Drivers of Local Alpha Diversity

Alpha diversity analysis showed that soil microbial richness in loams was significantly higher than in sands for all diversity indexes examined, except for the Simpson index for fungi (*p* = 0.51). The analysis also revealed distinct patterns in microbial richness and evenness across the studied sites and biotopes (Fig. 5, S1). For both bacteria and fungi, Chao1 and Shannon indices displayed graphical trends highly similar to the observed ASV richness. The Shannon index ranged from 4.5 to 6.5 for bacterial communities and from 2.5 to 4.2 for fungal communities. This visual consistency suggests a good robustness of the alpha-diversity patterns and points to a sufficient sequencing depth. Moreover, because the Shannon index reflects ASV richness and accounts for rare taxa, diversity changes may be largely driven by these rare taxa. To avoid redundancy, Chao1 and Shannon metrics are in the Supplementary Material (Fig. S1).

**Fig. 5.**
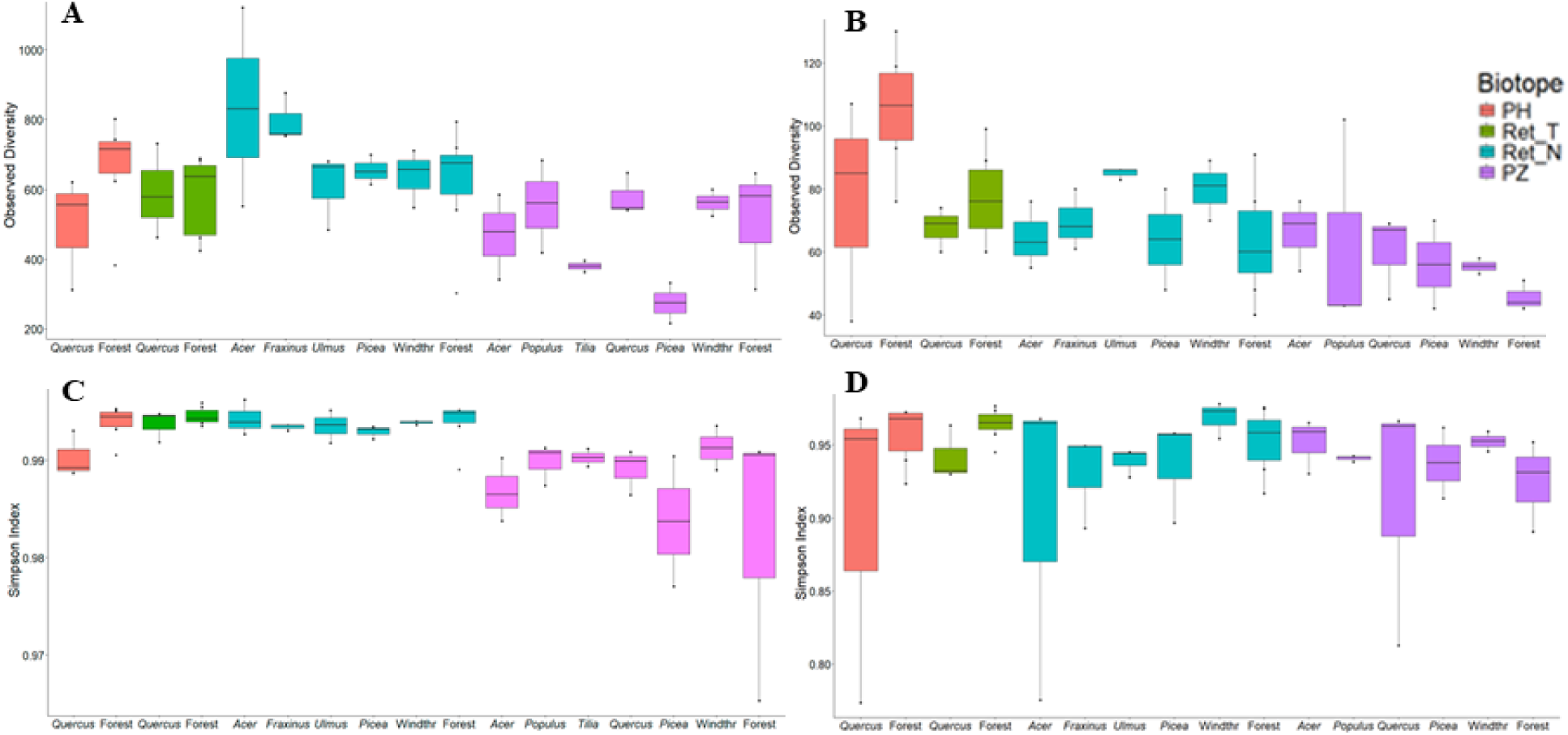
Alpha-diversity metrics across sites within biotopes. Bacterial (A, C) and fungal (B, D) observed richness (A, B) and inverse Simpson index (C, D). Values are calculated from ASVs.

ANOVA showed that biotopes explained 23% and 33% of the variation in observed richness for bacteria and fungi, respectively (*p* = 0.002 and 0.0001). Meanwhile, the study sites accounted for approximately 50% of the variance in diversity for both microbial groups (*p* = 0.006 in both cases). Tukey’s HSD test indicated that Ret_N biotope exhibits significantly higher bacterial ASV richness than the PZ biotope (*p* =0.001). Fungal ASV richness in the PH biotope was significantly higher than in both Ret_N and PZ biotopes (*p* = 0.002 and < 0.0001, respectively). Pairwise comparisons of specific sites revealed that bacterial diversity was significantly higher beneath *Acer* and *Fraxinus* deadwood in the Ret_N biotope compared with plots beneath *Picea* and *Tilia* deadwood in the PZ biotope (*p* < 0.05). Fungal diversity was significantly elevated in the Forest site in PH biotope compared to the background Forest sites in both Ret_N and PZ biotopes (*p* < 0.05). PZ biotope showed the lowest overall microbial richness and evenness.

The inverse Simpson index analysis revealed a high overall community evenness across most studied groups, with values consistently ranging between 0.90 and 0.99 (Fig. 5C, D). For the bacterial communities (Fig. 5C), the Simpson index remained remarkably stable and close to its maximum in the PH, Ret_T, and Ret_N biotopes, but exhibited a sharp decline and a high degree of dispersion within the PZ biotope sites (particularly beneath *Picea* CWD and in background Forest). This indicates a disruption of bacterial community evenness in podzols, likely driven by the dominance of specific taxa.

For fungal communities, the Simpson index (Fig. 5D) revealed substantial variations in community evenness, distinguishing it from the bacterial trends. While the Ret_T biotope maintained high and stable diversity metrics across all sites, the other biotopes were characterized by localized drops in the index. Notably, severe fluctuations and increased dominance of specific taxa (indicated by lower Simpson values around 0.77–0.81) were captured beneath *Quercus* CWD in PH and PZ biotopes, as well as beneath *Acer* CWD within the Ret_N biotope. Conversely, sites without logs (Forest and Windthr sites) consistently supported higher and more homogenous fungal evenness across the biotopes.

GLMs using negative binomial regression (GLM NB) revealed distinct, kingdom-specific responses of soil microbial alpha diversity to environmental and soil texture gradients (Fig. 6). Initial model selection based on the principle of parsimony demonstrated that a simple binary classification of soil texture explained variance in microbial richness (expressed as the number of ASVs) in relation to soil characteristics more effectively and with higher statistical significance than broad Biotope classifications including differences in soil types and windthrow events.

**Fig. 6.**
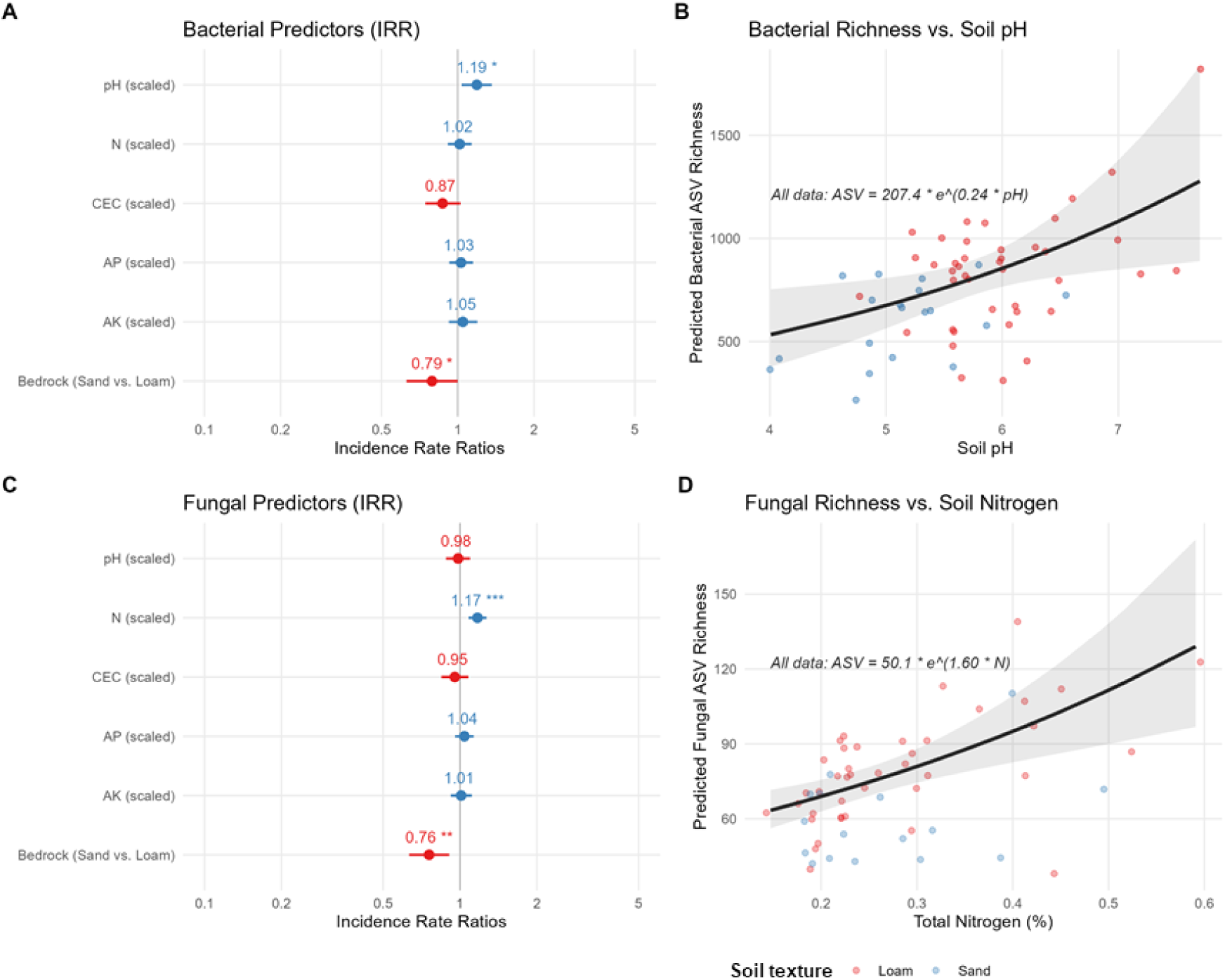
Generalized linear model (Negative Binomial) outputs for soil bacterial and fungal ASV richness. (A, C) Forest plot showing Incidence Rate Ratios (IRRs) with 95% confidence intervals for scaled bacterial and fungal community predictors. (B, D) Marginal effects plot displaying overall trend lines of predicted exponential increase in bacterial and fungal ASV richness along the unscaled chemical (pH and nitrogen) gradient. For plots A and C, predictors with intervals crossing the dashed line (1.0) are statistically non-significant; all continuous variables were standardized (scaled) prior to analysis. For plots B and D, individual sample points are color-coded by type of soil texture (Loam vs. Sand) to visualize data distribution, shaded areas represent 95% confidence intervals, and a single comprehensive regression equation is provided.

Bacterial ASV richness exhibited a highly significant, positive exponential relationship with soil pH (*beta* = 0.17 ± 0.07, *z* = 2.53, *p* = 0.0115) (Fig. 6A, B). For fungal ASV, total nitrogen (N) was identified as the dominant chemical predictor of alpha diversity (*beta* = 0.16 ± 0.04, *z* = 3.76, *p* = 0.00017) (Fig. 6C, D). In an alternative formulation, total carbon (C) exhibited a significant effect on fungal ASV richness (*p* = 0.001). Other chemical soil properties showed no significant effect on ASV richness. Beyond chemical characteristics, soil texture acted as a strong physical constraint: sandy soils significantly suppressed both bacterial and fungal diversity compared to loamy substrates (*beta* = -0.23 ± 0.12, *z* = -1.98, *p* = 0.05 for bacteria and *beta* = -0.28 ± 0.09, *z* = -2.98, *p* = 0.003 for fungi). Sandy soil significantly reduced bacterial and fungal ASV richness by approximately 21 and 24% compared to loams (*p* = 0.05 and 0.003, respectively).

### 3.5. Multivariate Constraints on Community Structure (beta-diversity)

#### 3.5.1. NMDS based on Bray–Curtis distances complemented by PERMANOVA

testing revealed distinct bacterial and fungal community separation across biotopes and sites (Fig. 7). Biotopes significantly affected beta diversity explaining 25% of the compositional variance for bacteria and 19% for fungi (both *p* = 0.001). For bacteria (stress = 0.123), ordination showed clear biotope segregation: PZ formed an isolated cluster along the positive NMDS1 axis, separate from PH and minimally overlapping with Ret_T and Ret_N. The latter two tightly clustered and overlapped centrally, whereas PH showed the widest dispersion, reflecting high intra-group variance. Despite a higher stress value (0.222), the ITS dataset preserved a similar trend, showing a directional shift of PZ toward positive NMDS1 values, though with greater spatial overlap among the Ret_T, PH, and Ret_N ellipses.

**Fig. 7.**
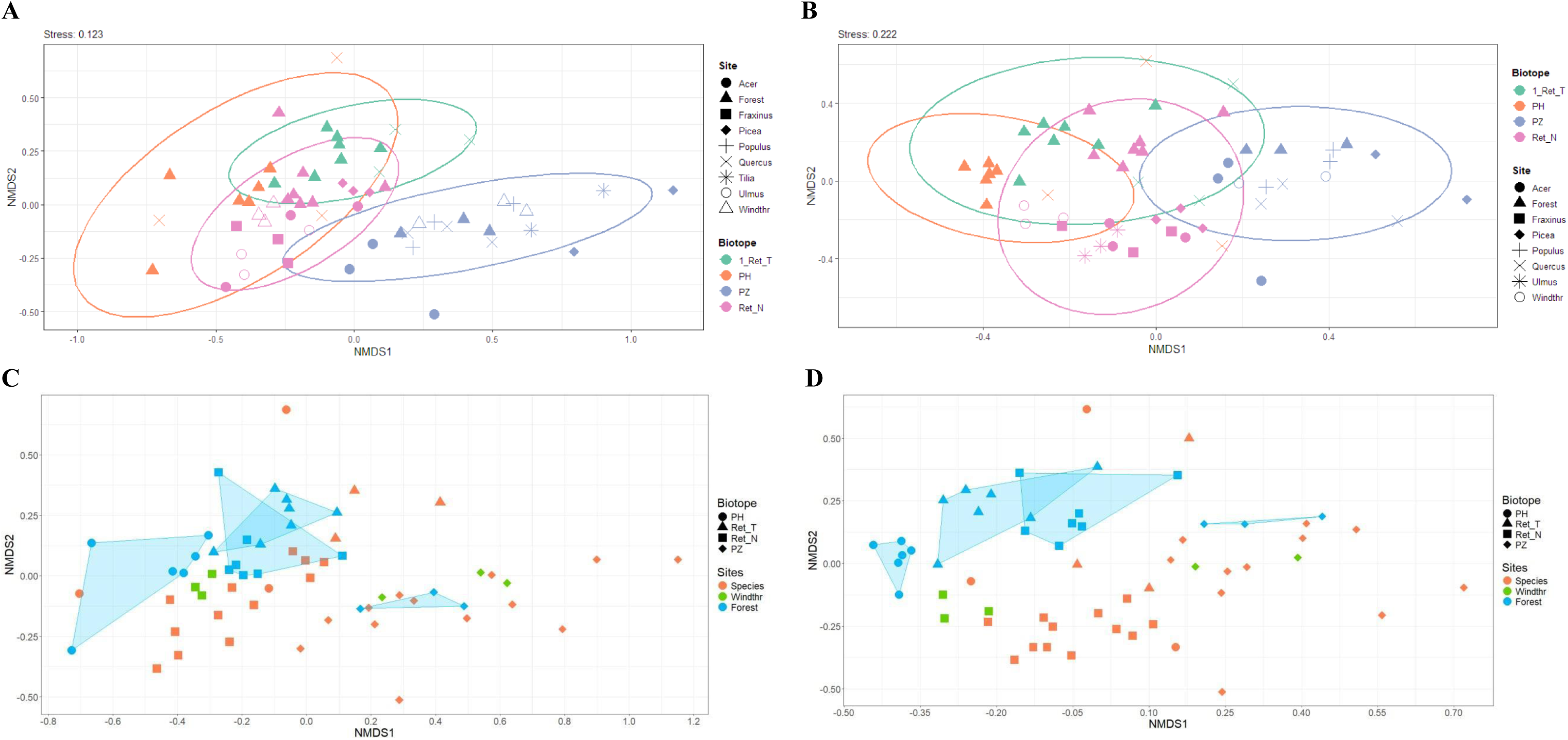
Similarity of bacterial (A, C) and fungal (B, D) community structures at the ASV level, visualized by NMDS ordination based on the Bray–Curtis distances (stress 0.123 and 0.222, respectively). Points represent individual samples with colors indicating the four distinct biotopes and shapes designating the nine sampling sites. A and B: continuous lines represent 95% confidence ellipses calculated for each biotope. C and D: convex hulls are shown for the log-free “Forest” control sites within each biotope.

Overall, wood decomposition substantially transformed microbial communities and increased beta-diversity, with log-free “Forest” controls clustering distinctly from log-influenced sites, especially for fungi (Fig. 7C, D). PERMANOVA confirmed that CWD tree species effects varied by biotope and microbial group. Within biotopes, bacterial structure in Forest sites differed significantly from log-influenced plots in Ret_N and Ret_T (*p* ≤ 0.009), while fungal Forest sites differed from log plots across all biotopes except Ret_T. Furthermore, CWD species composition significantly affected both communities within PZ (*p* ≤ 0.035), where *Picea* (both groups) and *Tilia* (bacteria) induced the strongest divergence from the control. In Ret_N, this effect was significant only for bacteria (*p* = 0.047; fungi: *p* = 0.221). Notably, the lack of significant bacterial differentiation of the Forest site in PZ, despite its visual NMDS segregation, stems from the high dimensionality of the V4 dataset (∼19,000 ASVs), which reduces PERMANOVA power at low sample sizes (*n* = 3); conversely, the more condensed ITS matrix (∼1,500 ASVs) successfully captured this habitat signal, despite lacking *Tilia* sequence reads.

#### 3.5.2. Distance-based Redundancy Analysis (dbRDA)

based on Bray-Curtis distances revealed a divergence in the biogeochemical architecture structuring soil bacterial and fungal community compositions across the forest floor (Fig. 8).

**Fig. 8.**
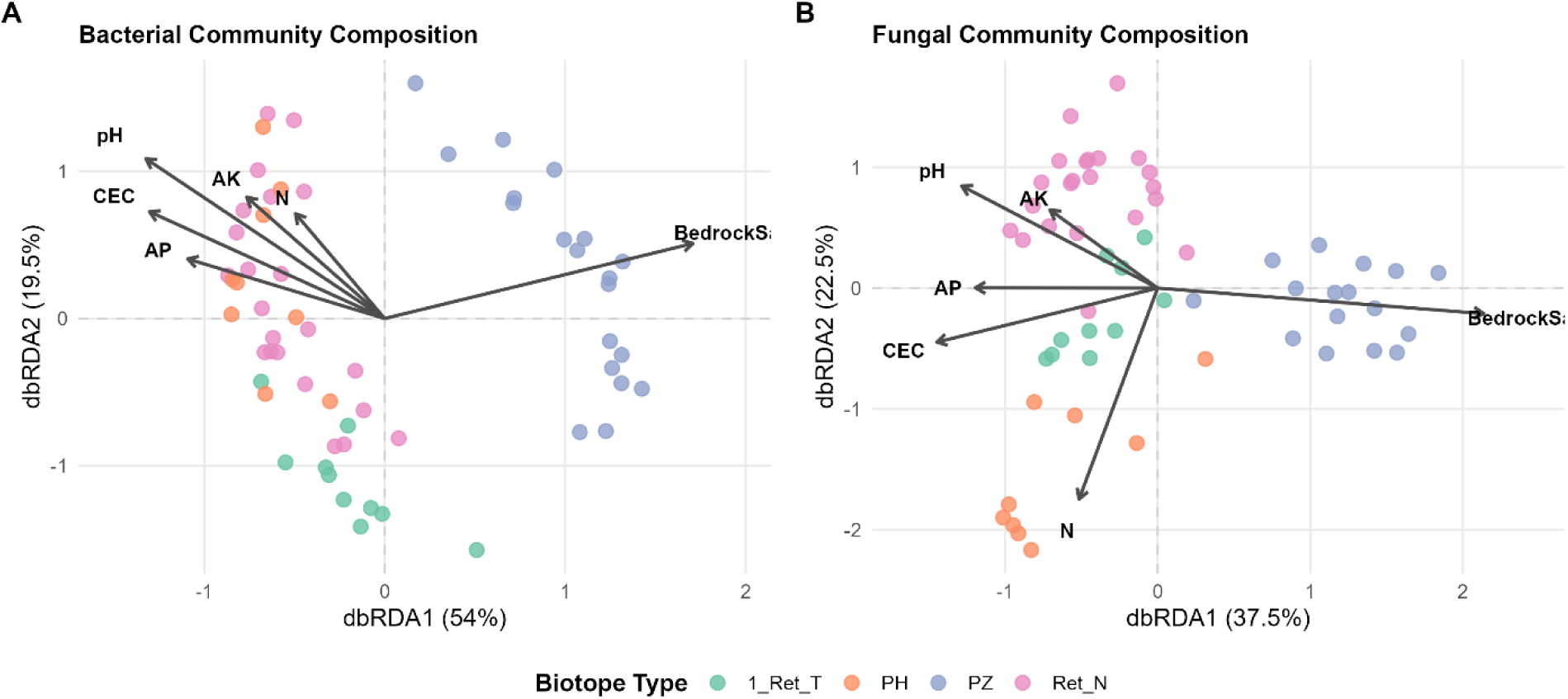
Distance-based Redundancy Analysis (dbRDA) ordination plots of soil bacterial (A) and fungal (B) community structures based on Bray-Curtis distances. Points represent individual soil samples color-coded by biotope types. Vectors indicate the direction and strength of soil environmental variables; significant predictors (*p* < 0.05) include soil texture (BedrockS), pH, total nitrogen (N) for both microbial groups, and availavle potassium (AK) exclusively for bacteria, whereas CEC and AP showed no significant or only marginal effects (see main text for the details). The percentages on the axes represent the proportion of constrained variance explained by each dbRDA axis.

In line with the alpha diversity trends, soil texture (*F* = 6.4, *p* = 0.001) and soil pH (*F* = 3.8, *p* = 0.001) emerged as the most powerful drivers structuring the bacterial communities. dbRDA also revealed that soil nutrient pool (total nitrogen N and available potassium AK), while having no effect on ASV richness, significantly influenced bacterial community composition (*F* = 1.6, *p* = 0.04 for N; *F* = 1.7, *p* = 0.02 for AK). This indicates that nitrogen and potassium gradients drive taxonomic turnover and species replacement rather than changes in net richness. CEC (*p* = 0.27) and AP (*p* = 0.53) exerted no significant influence on bacterial composition. These environmental pressures resulted in a distinct clustering pattern (Fig. 8A), where samples from the PH and Ret_N biotopes exhibited extensive overlap in the upper-left quadrant, driven by pH and nutrient vectors. In contrast, the Ret_T samples formed a completely distinct cluster isolated along the negative dbRDA2 axis, reflecting its distinct relationship with the measured soil properties.

Similar patterns were observed for fungal communities. Correspondingly, soil texture (*F* = 3.7, *p* = 0.001) and total nitrogen N (*F* = 2.05, *p* = 0.004) constrained fungal community composition, demonstrating that physical filtering and nitrogen gradients regulate both the richness and taxonomic identity of soil fungi. Soil pH acted as an additional powerful constraint (*F* = 2.02, *p* = 0.001); much like nutrient effects in bacteria, pH did not limit the net number of fungal ASVs but drove a taxonomic turnover along its gradient. CEC (p = 0.091) and AP (p = 0.081) showed only marginal trends, while AK (p = 0.231) had no detectable effect on fungal community shifts. Notably, the orthogonal effect of pH and N on fungal community composition indicates that these two environmental drivers act independently, each shifting the community structure along distinct ecological axes. This independence was clearly reflected in the contrasting fungal clustering (Fig. 8B), where Ret_T samples shifted closer to the PH biotope. Meanwhile, the Ret_N biotope formed a separate cluster that stretched along the negative dbRDA2 axis, aligned directly with the strong downward vector of total nitrogen (N), confirming that nitrogen availability serves as a critical, divergent filter for the soil mycobiome.

### 3.6. Bacterial and Fungal Gamma Diversity Profiles

Analysis of regional (gamma) diversity using iNEXT curves (Fig. 9) revealed contrasting accumulation trajectories between bacterial and fungal communities across the four biotopes, demonstrating a tight coupling with soil texture, microbial biomass (qPCR), and key environmental gradients identified via dbRDA.

**Fig. 9.**
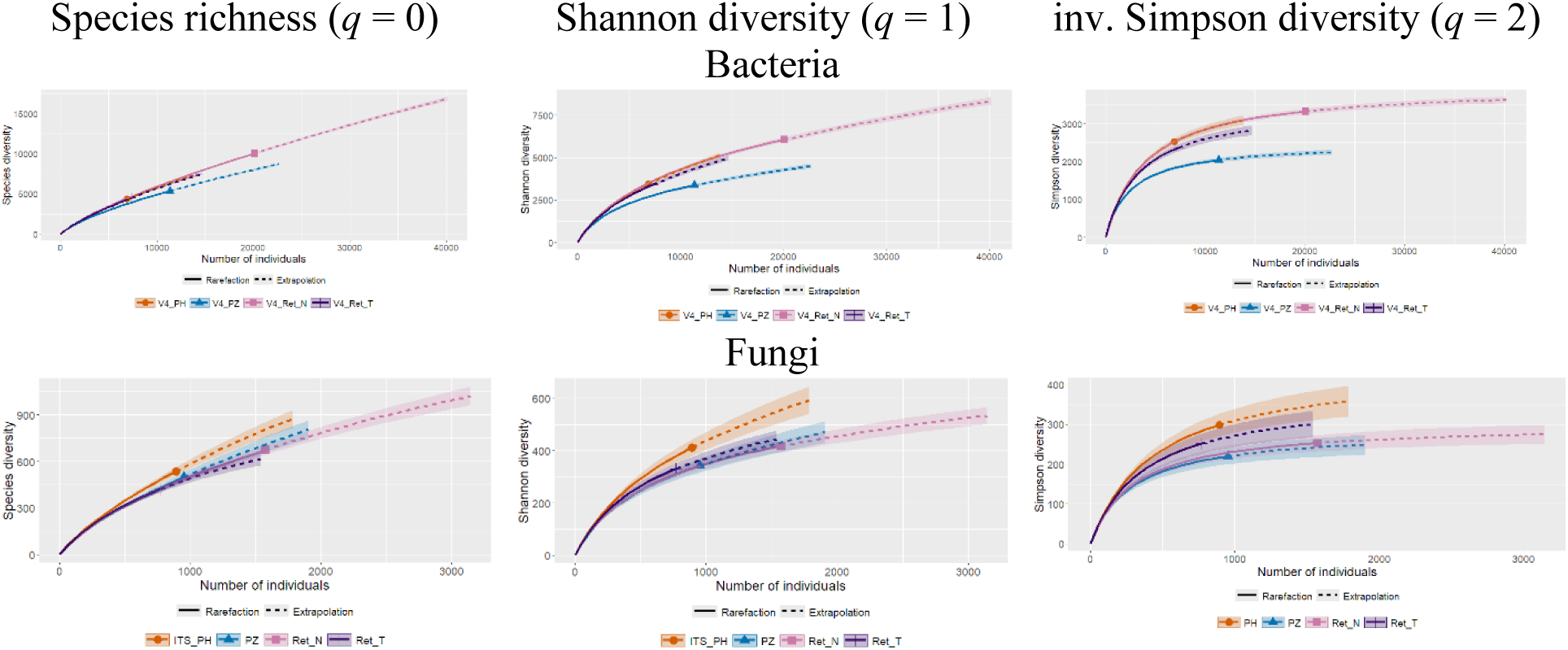
Sample-size-based interpolation (rarefaction) and extrapolation curves for soil bacterial (top) and fungal (bottom) gamma-diversity across four distinct biotopes based on Hill numbers: *q* = 0, species richness; *q* = 1, Shannon diversity; *q* = 2, inverse Simpson diversity. Solid lines represent rarefaction (interpolated) data, dashed lines represent extrapolation, and shaded areas denote 95% confidence intervals. Points mark the actual reference sample sizes.

For bacteria, the sandy PZ biotope consistently exhibited the lowest gamma-diversity among all studied biotopes across all Hill numbers (Fig. 9, top), matching its suppressed bacterial DNA load compared to loamy soils (*p* < 0.001). According to dbRDA and GLM regression, community structure and alpha-diversity were driven by soil texture, total N, AK, and primary pH as a master regulator. The smaller pool of ASVs in Ret_T, contrasted with the overlapping, maximum-richness trajectories of PH and Ret_N, demonstrates how these soil properties (specifically the highest values of pH and AK, Table 2) filter the bacterial community at a regional scale.

Fungi displayed a more complex, index-dependent pattern (Fig. 9, bottom) reflecting macroscale community evenness. Fungal community structure was driven primarily by soil texture, N, and pH (dbRDA), with nitrogen/carbon identified as the critical predictor for richness (GLM). Across all Hill numbers, the PH biotope maintained the highest, most stable fungal gamma-diversity, indicating a structurally balanced regional mycobiome. Interestingly, for species richness (*q* = 0), the fungal pool size in the sandy PZ biotope surpassed Ret_N and Ret_T, matching its significantly enhanced fungal DNA load (*p* < 0.001) which potentially facilitates a pool of low-abundance populations. However, at higher Hill numbers, structural divergence emerged: for Shannon diversity (*q* = 1), PZ, Ret_N, and Ret_T dropped below PH and became indistinguishable, whereas for the inverse Simpson index (*q* = 2), Ret_T approached PH, leaving PZ and Ret_N overlapping at lower asymptotic values. This shift across Hill numbers reveals distinct structural features of the regional mycobiomes. In the sandy PZ biotope, high fungal abundance supports a vast reservoir of rare ASVs (*q* = 0), but its core structure is shaped by localized drops in evenness and increased dominance of specific taxa (*q* = 2). Conversely, the upward shift of the Ret_T toward PH at *q* = 2 mirrors its high alpha-diversity evenness (Fig. 5) and aligns with the dbRDA clustering (Fig. 8): along the evaluated environmental gradients the Ret_T fungal community is structurally closer to PH than Ret_N is.

To further dissect the biotope-level trends, we evaluated the microbial gamma-diversity accumulation curves at the site level across the studied biotopes (Fig. 10).

**Fig. 10.**
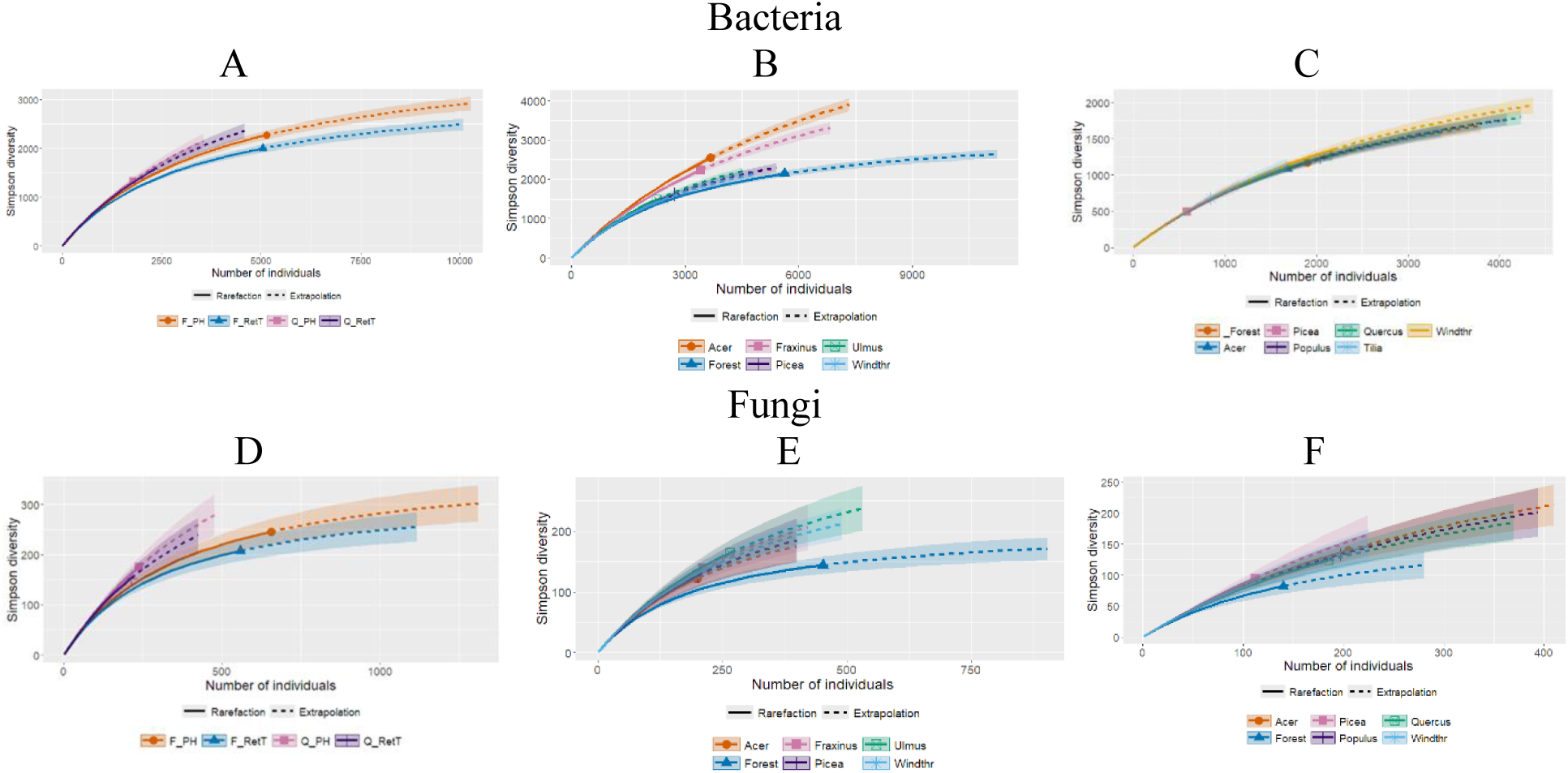
Sample-size-based interpolation (rarefaction) and extrapolation curves of Simpson diversity (*q* = 2) in bacterial (up) and fungal (bottom) communities at the site level within biotopes. A and D: PH and Ret_T; B and E: Ret_N; C and F PZ biotopes. Solid lines represent rarefaction (interpolated) data, dashed lines represent extrapolation, and shaded areas denote 95% confidence intervals. Points mark the actual reference sample sizes.

In the old-growth forest with isolated windthrows (PH and Ret_T biotopes), *Quercus* CWD acted as a unifying ecological factor standardizing bacterial and fungal community structures (Fig. 10A, D; Fig. S2). The accumulation curves for the sites beneath *Quercus* logs in both PH and Ret_T biotopes completely overlapped, revealing a significantly higher pool of ASVs compared to their respective log-free background forest controls (apart from *q* = 0 for bacteria, where all curves overlapped completely). The log-free “Forest” sites generally maintained their landscape-specific signatures: microbial communities in PH (the biotope with supreme levels of C and N, Table 2) were richer than those in Ret_T (although this difference was statistically significant only for dominant bacteria; *q* = 2). Notably, the soil beneath the oak CWD exhibited greater gamma diversity of the microbiome compared with the forest background, although the mean and median numbers of microbial ASVs per sample dropped under the logs (Fig. 5).

In the loamy mass-windthrow Ret_N biotope, the microbial gamma-diversity revealed a highly partitioned structure shaped by the windthrow disturbance mosaic, showing contrasting successional responses between the bacterial and fungal communities (Fig. 10B, E; S3). For bacteria, (Fig. 10B, S3), the impact was heavily localized: the soil environments beneath *Acer* and *Fraxinus* CWD drastically outpaced the others, breaking away to form an exceptionally high pool of ASVs. In contrast, the remaining windthrow-affected sites, including the other CWD inputs (*Ulmus, Picea*) and the log-free windthrow gaps (Windthr), clustered tightly with the undisturbed background forest floor (Forest), which maintained a comparably high baseline across all diversity indexes. Conversely, the fungal community (Fig. 10E, S3) exhibited a fundamentally different structural pattern. Instead of isolated single-site peaks, fungal gamma-diversity was characterized by a tight, elevated cluster of almost all windthrow-associated microhabitats, including the log-free Windthr gaps, which shared overlapping 95% confidence intervals across all Hill numbers. The undisturbed forest floor was the only site that failed to shift, separating from the entire windthrow mosaic and plateauing at a baseline lower than the newly established fungal cluster. This indicates that on loamy soils, the windthrow event triggers a highly site-specific hot-spot dynamic for bacterial richness centered on select log types, leaving the rest of the mosaic at the high baseline of the background forest, whereas it drives a uniform, widespread expansion of fungal diversity across the entire disturbed landscape.

In the sandy PZ biotope (Fig, 10C, F), the microbial gamma-diversity was also shaped by the windthrow-induced disturbance mosaic and the response varied fundamentally between bacteria and fungi (Fig. S4). The bacterial community showed a highly homogenous pattern in species richness (*q* = 0), with all curves overlapping. However, Shannon diversity (*q* = 1) revealed a distinct shift, where the undisturbed background forest floor ran visibly below all windthrow-affected sites, including both the CWD inputs and the log-free windthrow gaps (S4). This divergence exclusively under *q* = 1 indicates that while the total bacterial species pool remains uniform across the biotope, the bacterial community structure in the background forest is skewed with rare ASV accounting for a smaller fraction of the total abundance (Fig. 5C, S1). The overall windthrow disturbance structurally alters the balance, promoting a more even distribution of bacterial abundances rather than expanding the absolute richness. In turn, the fungal community revealed a more stratified structure, showing a clear division between the transformed windthrow microhabitats and the background forest. While the curves for all windthrow-associated sites (*Acer, Picea, Quercus, Populus*, and the log-free Windthr) tightly clustered together along an elevated diversity trajectory, the background forest floor formed a distinctly lower, isolated baseline across all three Hill numbers. This indicates that while the windthrow event (through both deadwood deposition and the creation of open gaps) uniformly enhances and differentiates soil fungal diversity from the surrounding matrix, its impact on the bacterial community is restricted to restructuring taxon evenness.

## General Conclusion / Synthesis

In conclusion, our cross-biotope synthesis demonstrates that the impact of windthrow disturbances on soil microbial gamma-diversity is determined by a complex interplay between the scale of the disturbance, host tree species, and underlying soil physico-chemical matrices. While deadwood deposition consistently acts as a qualitative catalyst for expanding the total species pool, the structural pathways of microbial species accumulation are modulated by distinct responses of bacterial and fungal communities. This is supported by background quantitative PCR trends, which showed a general suppression of bacterial abundance beneath the logs as a spatial position effect, whereas fungal DNA abundance was highly context-dependent, filtered by a significant interaction between biotope and CWD tree species identity. Despite these localized shifts in biomass, the overall landscape response highlights a profound spatial decoupling between local niche saturation and gamma-level species accumulation, shifting dynamically across different forest disturbance regimes.

In landscapes shaped by catastrophic, large-scale windthrows (Ret_N and PZ), the microbial response is highly domain-specific and heavily filtered by baseline soil properties. On the loamy soils of the Ret_N biotope, which are characterized by a significantly higher baseline pH and available potassium but standard carbon and nitrogen levels (Table 2), the specific micro-environmental conditions developing beneath *Acer* and *Fraxinus* CWD act as a potent driver, sharply increasing bacterial diversity and forming highly localized richness hot-spots. In turn, on the sandy soils of the PZ biotope, where the sand matrix systemically suppresses bacterial biomass, the overall windthrow disturbance acts as a stabilizing force that restructures community evenness across the entire mosaic (encompassing both logs and open gaps) rather than expanding the absolute species pool. Fungal communities in these mass-windthrow areas exhibit a uniform systemic shift, expanding their overall species richness relative to the background forest regardless of soil texture and driven by resource availability (nitrogen / carbon) and soil pH.

A fundamentally different ecological mechanism operates in old-growth forests characterized by isolated, small-scale windthrows (PH and Ret_T). Here, large oak (*Quercus*) logs act as a powerful “ecological standardizer” erasing baseline differences between carbon- and nitrogen-rich Phaozems and nutrient-depleted Retisols. While undisturbed forest floors keep distinct, soil-dependent profiles with Phaeozems hosting richer communities than Retisols, the addition of oak CWD unifies microbial trajectories across both soil types. This homogenizes community structures and boosts regional gamma-diversity. Crucially, while oak logs enhance the overall landscape pool of ASVs (gamma-diversity), they simultaneously trigger a localized reduction in alpha-diversity per individual sample, highlighting a spatial decoupling between local niche saturation and regional species accumulation. Collectively, these findings reveal that while large-scale windthrow events drive diverse, edaphically-filtered successional pathways across open mosaics, isolated old-growth deadwood functions as a uniform ecological catalyst, creating consistent microbial refugia that enrich forest soil diversity at the landscape scale.

## 4. Discussion

### 4.1. Edaphic and Structural Constraints of the Soil Matrix on Microbial Abundance and Diversity

The baseline physical structure of the soil acts as a primary ecological filter, regulating microbial communities at the local and regional scales. Our study reveals that the soil texture determines microbial DNA content, local alpha-diversity (niche availability), which subsequently scales up to define the regional species pool (gamma-diversity).

The systemic suppression of local bacterial DNA content (qPCR) and bacterial alpha-diversity in the sandy soil has been widely demonstrated (e.g., Girvan et al., 2003; Hemkemeyer et al., 2018), highlighting the severe physical constraints imposed by coarse-textured profiles (Hemkemeyer et al., 2018, Seaton et al., 2020). In contrast to loamy matrices, sandy environments possess a significantly lower specific surface area and lack the capacity to form stable organo-mineral complexes (Schimel et al., 1994; Sollins et al., 1996; von Lützow et al., 2006; De Vries et al., 2012). In loamy soils, these clay-humus sheaths physically protect microbial cells and extracellular bacterial DNA from enzymatic degradation and hydrological leaching (Schmidt et al., 2011; Nannipieri et al., 2012). In the macroporous sandy matrix of the PZ biotope, the lack of such protective micro-niches likely imposes a strict physical limit on local carrying capacity. This structural deficit keeps bacterial biomass, alpha-diversity (sample richness), and gamma-diversity (the total species pool) low, regardless of carbon and nitrogen availability.

*Alphaproteobacteria,* the dominant group in the studied forests, comprise diverse bacterial taxa crucial for carbon, nitrogen, and sulfur cycling (Kersters et al., 2006). Predominantly chemoheterotrophic, these bacteria frequently form symbiotic relationships with plants and contribute to nitrogen fixation, showing a strong association with the forest rhizosphere (Lladó et al., 2017). *Verrucomicrobia,* which shared dominance in the soils, are globally ubiquitous, though their specific ecological roles remain partially elusive (Lladó et al., 2017; Meena et al., 2023). Nevertheless, their primary function in soil environments involves the decomposition of plant-derived organic matter (Rakitin et al., 2024).

The structural and physiological difference between the two microbial domains (bacteria and fungi) becomes evident when analyzing the fungal response within the same sandy matrix. Although the sand matrix did not increase local fungal alpha-diversity (which, except for the Simpson index, was significantly lower than in loams), fungal DNA content was markedly enhanced. This suggests that while the macro-porous environment limits local alpha-diversity, its high fungal biomass supports the diffuse persistence of filamentous growth forms. Fungal hyphae can bridge physical gaps between sand pores to translocate moisture and nutrients from localized organic matter hot-spots (Allen, 2007; Guhr et al., 2015). When scaled up to the landscape level, this broad biomass foundation allows a reservoir of rare, low-abundance populations to persist, facilitating a decoupling where a standard local alpha-diversity profile scales up to an elevated regional species richness trajectory on iNEXT curves (Fig. 9). Under higher Hill numbers (*q* = 2), the subsequent decrease in fungal diversity in the sandy PZ compared to loamy biotopes tightly reflects the local drops in evenness (Fig. 5D). This combination of high fungal DNA content and low alpha-diversity indicates intense competitive exclusion, where local micro-habitats are dominated by a small number of highly specialized, competitively superior taxa outcompeting others under these specific conditions.

The physical properties of coarse sand impose severe desiccation and temperature fluctuations, favoring the highly resilient, melanized ectomycorrhizal ascomycete *Cenococcum geophilum* (Jany et al., 2003; Fernandez & Koide, 2013; Peter et al., 2016). Concurrently, nitrogen-limited and carbon-rich micro-habitats within the matrix are often monopolized by the oligotrophic, psychrotolerant, and cellulolytic saprotrophs *Geomyces, Pseudogymnoascus roseus*, and *P. appendiculatus* (Rice, Currah, 2006; Lorch et al., 2013). The physiological and metabolic specialization of these taxa allows them to outcompete generalist decomposers and generate an immense biomass footprint, structurally homogenizing local micro-habitats via competitive exclusion. In general, there are many mycorrhizal fungi on sandy soils, such as *Cenococcum*, *Oidiodendron* and some *Helotiales*.

### 4.2. Localized Deadwood Effects and Chemical Footprints of Tree Species on Soil Microbial Communities

The deposition of CWD triggers a profound spatial restructuring of the forest floor, shifting microbial assembly from macro-environmental edaphic control to localized, micro-habitat-driven successional pathways. At the scale of the individual deadwood footprint, decomposing logs operate as a severe quantitative and local qualitative filter, significantly suppressing bacterial biomass and altering microbial diversity. Significant reduction in bacterial DNA content in the soil beneath CWD, compared to log-free sites, confirms the suppressive effect on microbial activity under the logs that we previously observed in loamy soils using the substrate-induced respiration method (Khanina et al., 2024; Smirnov et al., 2024). These findings point to a robust spatial effect that goes beyond the general landscape types.

Decomposing logs release massive fluxes of labile organic carbon, but they concurrently inject high concentrations of antimicrobial secondary metabolites into the underlying soil solution, such as polyphenols, tannins, and organic acids (Stutz et al., 2017; Bärlocher et al., 2017). Furthermore, fungal wood-degraders extensively colonizing the logs drastically lower the pH of the substrate through the secretion of organic acids and release reactive oxygen species during cellular metabolism, creating highly stressful conditions for standard soil bacteria (de Boer & van der Wal, 2008; Rousk et al., 2010). We have also confirmed a decrease in pH in wood of seven broad-leaved trees and spruce at intermediate stages of wood decomposition at our study site (Khanina et al., 2023). Additionally, the physical mass of a large log acts as a physical barrier that limits light penetration, moderates temperature fluctuations, and severely alters gas and moisture dynamics (Stokland et al., 2012; Khanina et al., 2024). This combination of chemical toxicity, active fungal antagonism, and an altered microclimate creates a highly stressfull microenvironment in which highly specialized, acid-tolerant, or oligotrophic taxa gain a competitive advantage (Kielak et al., 2016; Stutz et al., 2017; Tláskal et al., 2017; Moll et al., 2018; Tláskal, Baldrian, 2021; Steinebrunner 2025). However, the effects on the soil depend significantly on the species of the fallen tree.

On loamy soils in the Ret_N biotope, *Acer* and *Fraxinus* CWD act as local chemical stimulators of bacterial diversity. Both of these species are well-studied ‘soil-improving’ tree species, characterised by wood with a high alkali content, which releases significant fluxes of alkaline cations and maintains a higher pH level compared with other species (Augusto et al., 2002; Muhr et al., 2016; Khanina et al., 2023). Specific bacterial communities, in which *Acidobacteria* are absent, formed beneath *Acer* CWD. In contrast, the proportion of *Acidobacteria*, which are commonly found under CWD (Wang et al., 2023? 2024; Błońska et al., 2024), increased in our study in loams specifically beneath *Quercus* and *Picea* CWD, which are characterised by particularly low pH values (3.5 and 4.1, respectively) at intermediate stages of decomposition (Khanin et al., 2023), leading to a reduction in soil pH in loams beneath the CWD of these species (Khanina et al., 2024). A striking feature of the dominant fungal communities beneath ash tree trunks is the high proportion of unidentified *Inocybaceae*. As the ash tree (*Fraxinus*) itself does not form ectomycorrhizae, the peak in *Inocybaceae* under *Fraxinus* CWD (in Ret_N) may be explained by the fact that windfall alters the physical properties of the soil and may provide shelter for other species. Importantly, as ash windfall decomposes, it releases a large amount of dissolved carbon (Lasota et al., 2022), which can be utilised.

As supported by our dbRDA alignment, the rapid release of available potassium (AK) and the localized maintenance of a favorable, near-neutral pH under these specific logs alleviate environmental stress, triggering a localized explosion of specialized bacterial richness (alpha-diversity hot-spots) without duplicating the background soil signature. *Acer* in PZ supports relatively high alpha and gamma-diversity. Within the loamy Ret_N biotope, high values of fungal diversity were observed beneath the *Ulmus* CWD and in the log-free area. Meanwhile, within the fungal community beneath *Ulmus* CWD, the Ascomycete genus *Lasiosphaeris* is present amongst the dominant species; this bioindicator points to active wood-degradation processes and the final stages of lignin and cellulose decomposition, as well as favorable hydrothermal conditions (Põlme et al., 2020). *Picea* on loamy soils maintains a moderate level of diversity. In the PZ biotope, spruce exhibits the lowest alpha diversity indices for bacteria, alongside the highest bacterial DNA loads. The proportion of *Acidobacteria* increases in loamy soils beneath *Picea* CWD, which are characterised by particularly low pH values (4.1) at intermediate stages of decomposition (Khanina et al., 2023).

A completely inverse chemical strategy is exhibited by large *Quercus* logs in the PH and Ret_T biotopes. Decomposing oak wood releases intense fluxes of highly recalcitrant lignin, complex polyphenols, and hydrolyzable tannins, creating an exceptionally rigorous and hostile chemical filter (Stokland et al., 2012). This aggressive, uniform chemical matrix effectively overrides the baseline shifts in soil pH and AK that otherwise differentiate the background soil profiles. The oak logs standardize the soil as bacterial, as fungal community structures beneath the logs, pulling the communities from two fundamentally different soil types (PH and Ret_T) into a single, tightly clustered multidimensional space. In PH, the dominance of active endophytes (*Leptodontidium*, *Helotiales*) and stable decomposers (*Humicola*) may indicate a stable food supply and minimal stress. In Ret_T, there is excessive moisture, whilst humidity is less stable; hydrophilic yeasts *Satiozyma*, black yeasts *Cyphellophoraceae* and moss ascomycetes *Hyaloscyphaceae* dominate. In the sandy PZ, *Quercus* logs support relatively high levels of alpha and gamma bacterial and fungal diversity and high beta-diversity, especially in the fungal community. Here stress-tolerant *Pseudogymnoascus* and the specialised saprophyte *Leptodontidium* dominated, reflecting an abundance of hard-to-degrade organic matter (cellulose and lignin) and acidic conditions, prevailed.

Our iNEXT and NMDS analyses reveal that within all biotopes decaying logs function as overriding ecosystem hotspots that bypass macro-chemical and soil texture constraints. Instead of shifting the microbial community in a uniform direction, CWD acts as a micro-environmental heterogeneity generator. By introducing highly localized, contrasting conditions, such as patches of organic acids, variable moisture retention, and concentrated inputs of wood-derived labile carbon, CWD induces an exponential expansion of beta diversity (high spatial turnover). Maximum values of gamma-diversity, whilst alpha-diversity values are not always high and are sometimes quite low, underscores that while soil chemistry and texture dictate the baseline matrix of the forest floor, the cumulative biodiversity of the forest ecosystem is critically sustained by localized, stochastic micro-habitats generated by decaying woody debris.

### 4.3. Disturbance Regimes: Mass Windthrows vs. Isolated Treefalls

When scaled up from individual log footprints to the landscape level, the spatial architecture of forest disturbances emerges as a major determinant of microbial gamma-diversity trajectories. Our findings reveal a fundamental difference between the successional pathways triggered by catastrophic, large-scale mass windthrows (Ret_N and PZ) versus those driven by isolated, small-scale treefalls in old-growth stands (PH and Ret_T).

In landscapes shaped by mass windthrows, the microbial response is governed by a heterogeneous mosaic of open spaces and highly variable deadwood inputs. The removal of the canopy shield across vast areas fundamentally alters the macroscale microclimate, introducing severe temperature oscillations, altering topsoil moisture cycles, and accelerating organic matter turnover in the log-free windthrow gaps (Bunnell et al., 2002; Stokland et al., 2012). For the bacterial community, this open mosaic acts as a systemic stabilizing force rather than a creator of richness hot-spots, particularly in sandy matrices (PZ) where texture limits biomass. The mass disturbance structurally shifts community assembly by altering taxon evenness across the entire landscape, balancing the dominance of core background taxa.

For the fungal domain, however, this mass-windthrow mosaic drives a uniform, collective expansion of overall gamma-diversity (*q* = 0) that completely outpaces the undisturbed forest floor, operating independently of the underlying soil texture. Because heterotrophic fungi are primary wood-degraders equipped with complex enzymatic machinery to break down recalcitrant polymers, their successional trajectories are heavily governed by resource availability (nitrogen and carbon) and soil pH rather than the physical structure of the mineral matrix (de Boer & van der Wal, 2008; Tláskal et al., 2017). Supported by the continuous, high-quality organic matter inputs accumulated over a century in these broadleaf stands, fungal hyphae efficiently bridge the spatial gaps between CWD inputs and the open Windthr zones, establishing a widespread, highly connected landscape-scale mycobiome.

A fundamentally different ecological mechanism operates under the isolated treefall regime of old-growth forests. Here, the undisturbed forest matrix remains dominant, maintaining distinct, soil-dependent baseline profiles where the carbon- and nitrogen-rich Phaeozems (PH) support richer communities than the depleted Retisols (Ret_T). Within this stable matrix, isolated oak logs act as a potent “ecological standardizer” (Stokland, 2012). By overriding landscape-specific soil signatures, the deadwood unifies the microbial successional trajectories across different soil types. Crucially, while the intense chemical filtering of oak deadwood suppresses individual sample abundance (qPCR) and depresses mean local alpha-diversity, the high compositional uniqueness (beta-diversity) of these specialized sub-log communities acts as the primary engine for landscape-scale enrichment. When accumulated via iNEXT, these highly distinct “islands” introduce entirely new sets of ASVs to the regional pool, resulting in a sharp upward trajectory in total gamma-diversity. This reveals a profound spatial decoupling between local niche saturation and landscape species accumulation: catastrophic mass-windthrows generate diverse, edaphically-filtered successional matrices across open gaps, whereas isolated old-growth deadwood functions as a uniform ecological catalyst, establishing highly consistent microbial refugia that enrich forest soil diversity at the landscape scale.

Our findings demonstrate a profound spatial decoupling in forest floor microbial communities, where a localized reduction in alpha-diversity under coarse woody debris (CWD) drives a sharp increase in landscape-scale gamma-diversity. While standard soil physicochemical properties rigidly dictate baseline microbial richness (Fierer, Jackson, 2006), CWD functions as a powerful ecological catalyst and micro-environmental hotspot that overrides these baseline constraints. By filtering out dominant matrix-associated taxa, deadwood inputs create highly specialized micro-refugia with unique amplicon sequence variants (ASVs). This niche selection triggers pronounced compositional dispersion (NMDS/dbRDA) and high beta-diversity, which is particularly pronounced beneath oak (*Quercus*) logs. This aligns with recent fine-scale deadwood studies showing that localized microhabitats significantly inflate overall landscape heterogeneity by masking background soil signatures (Pioli et al., 2023; Zhu et al., 2026).

The structural manifestation of this spatial decoupling shifts dynamically depending on the forest disturbance regime. In landscapes recovering from mass-windthrow events, the microbial response is heavily filtered by the interaction between wood traits and baseline soil textures. On loamy soils within windthrow mosaics, this interaction triggers a highly partitioned, site-specific hotspot dynamic. While most disturbed microhabitats, including spruce, elm, and log-free gaps, remain anchored to the baseline profile of the background forest, the soil environments beneath *Acer* and *Fraxinus* CWD break away to form an exceptionally large, distinct pool of unique ASVs. Such host-specific breakaway dynamics support the “hotspots” framework of soil matrix partitioning (Kuzyakov, Blagodatskaya, 2015; Bosch et al., 2023), confirming that broad-scale disturbances like windthrows do not homogenize the floor if specific, chemically distinct tree species act as filters for rare or specialized microbial lineages (Žifčáková et al., 2016; Baldrian et al., 2023). Ultimately, iNEXT accumulation profiles confirm that CWD does not merely multiply common soil taxa, but enriches the forest landscape by injecting critical beta-level heterogeneity into the soil matrix.

## 5. Conclusions

Our study provides robust evidence of a profound spatial decoupling in forest floor microbial communities, demonstrating that coarse woody debris (CWD) acts as a powerful ecological catalyst rather than a mere multiplier of common soil taxa. While baseline soil physicochemical properties rigidly dictate local richness, individual deadwood inputs override these matrix constraints by filtering out dominant taxa and establishing highly specialized micro-refugia. This intense niche selection, particularly pronounced beneath *Quercus* CWD and further modulated by tree species-specific filters like *Acer* and *Fraxinus* within post-windthrow mosaics, fuels an exceptional landscape-scale beta-diversity enrichment. Ultimately, these findings underscore that the spatial heterogeneity injected by deadwood dynamics is a fundamental driver of regional taxonomic capacity (gamma diversity), implying that retaining diverse CWD inputs is critical for buffering forest microbiomes against broad-scale environmental disturbances.

---

**Table S1.**
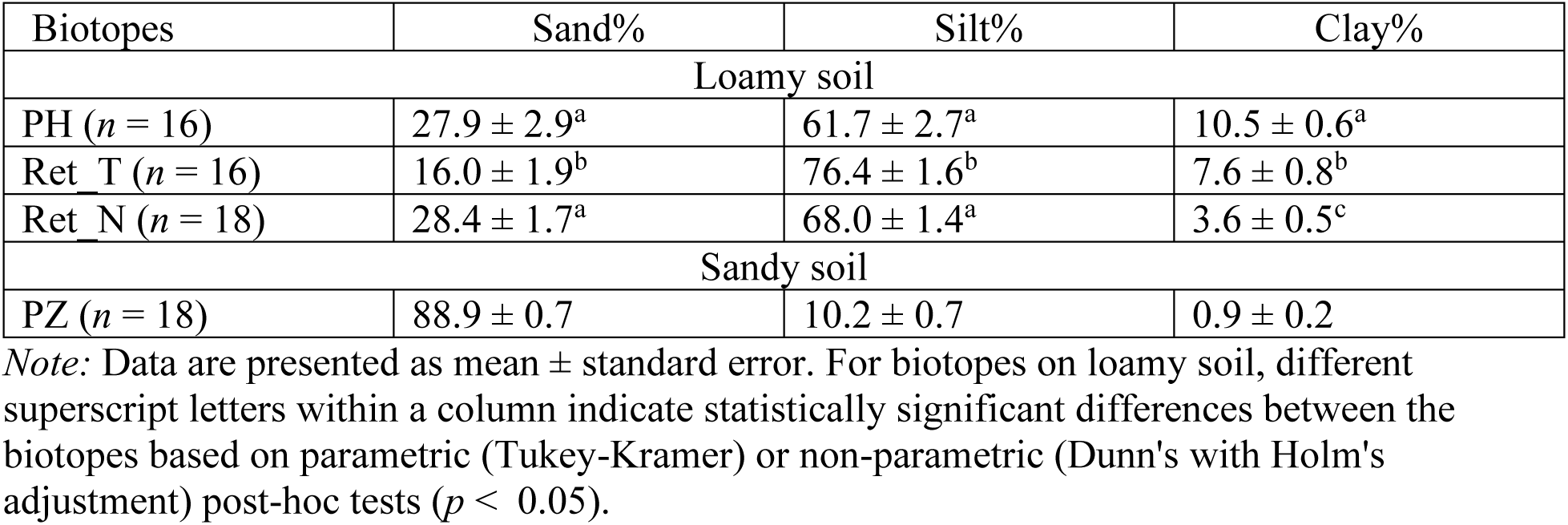
Soil texture in the studied biotopes (from Khanina, Bobrovsky, 2024 with modifications).

| Biotopes | Sand% | Silt% | Clay% |
| --- | --- | --- | --- |
| Loamy soil |  |  |  |
| PH ($n = 16$) | $27.9 \pm 2.9^a$ | $61.7 \pm 2.7^a$ | $10.5 \pm 0.6^a$ |
| Ret_T ($n = 16$) | $16.0 \pm 1.9^b$ | $76.4 \pm 1.6^b$ | $7.6 \pm 0.8^b$ |
| Ret_N ($n = 18$) | $28.4 \pm 1.7^a$ | $68.0 \pm 1.4^a$ | $3.6 \pm 0.5^c$ |
| Sandy soil |  |  |  |
| PZ ($n = 18$) | $88.9 \pm 0.7$ | $10.2 \pm 0.7$ | $0.9 \pm 0.2$ |

*Note:* Data are presented as mean ± standard error. For biotopes on loamy soil, different superscript letters within a column indicate statistically significant differences between the biotopes based on parametric (Tukey-Kramer) or non-parametric (Dunn's with Holm's adjustment) post-hoc tests ( $p < 0.05$ ).

## Notes

https://figshare.com/articles/dataset/_/30593849

### Competing Interest Statement

The authors have declared no competing interest.

## Literature

Abarenkov, Kessy; Zirk, Allan; Piirmann, Timo; Pöhönen, Raivo; Ivanov, Filipp; Nilsson, R. Henrik; Kõljalg, Urmas (2025): UNITE QIIME release for Fungi 2. UNITE Community. 10.15156/BIO/3301242

Allen, M. F. Mycorrhizal Fungi: Highways for Water and Nutrients in Arid Soils. Vadoze Zone Journal. 2007. 6(2): 291–297. doi: 10.2136/vzj2006.0068

Anderson M.J. 2017. Permutational multivariate analysis of variance (PERMANOVA). In Wiley StatsRef: Statistics Reference Online. Balakrishnan N., Colton T., Everitt B., Piegorsch W., Ruggeri F., Teugels J.L., Eds. Hoboken, NJ: John Wiley and Sons, Ltd., pp. 1–15. 10.1002/9781118445112.stat07841

Augusto, L., Ranger, J., Binkley, D., & Rothe, A. (2002). Impact of several common tree species of temperate forests on soil fertility and the properties of the soil profile. Annals of Forest Science, 59(3), 233–253.

Baldrian P., López-Mondéjar R., Kohout P. 2023. Forest microbiome and global change. Nat. Rev. Microbiol. 21 (8), 487–501. 10.1038/s41579-023-00876-4

Bani A., Pioli S., Ventura M., Panzacchi P., Borruso L., Tognetti R., Tonon G., Brusetti L. 2018. The role of microbial community in the decomposition of leaf litter and deadwood. Appl. Soil Ecol. 126, 75–84. 10.1016/j.apsoil.2018.02.017

Bardgett R., van der Putten W. 2014. Belowground biodiversity and ecosystem functioning. Nature. 515, 505–511. 10.1038/nature13855

Bärlocher, F., Gessner, M.O., Graça, M.A.S. (eds.) Methods to Study Litter Decomposition. A Practical Guide. Springer, Dordrecht. 2020. 10.1007/978-3-030-30515-4.

Błońska E., Ważny R., Górski A., Lasota J. 2024. Decomposing benefits: examining the impact of beech deadwood on soil properties and microbial diversity. Sci. Total Environ. 930, 172774. 10.1016/j.scitotenv.2024.172774

Bobrovsky, M.V. Forest Soils of European Russia: Biotic and Anthropogenic Factors of Formation (Tovarishchestvo Nauchnykh Izdanii KMK, Moscow, 2010) [in Russian].

Bobrovsky M.V., Stamenov. M.N., The catastrophic windfall of 2006 on the territory of the Kaluga Zaseki Nature Reserve, Lesovedenie, 2020, no. 6, pp. 523–536.

de Boer, W., van der Wal, A. (2008). Interactions between saprotrophic basidiomycetes and bacteria. In: British Mycological Society Symposia Series. Chap. 8. Eds: Lynne Boddy, J. C. F. van Pieter W. (Cambridge, MA: Academic Press) 28: 143–153.

Bolyen E., Rideout J.R., Dillon M.R. et al. 2019. Reproducible, interactive, scalable and extensible microbiome data science using QIIME 2. Nat. Biotechnol. 37 (8), 852–857. 10.1038/s41587-019-0209-9

Bosch, J., Némethová, E., Tláskal, V., Brabcová, V., & Baldrian, P. (2023). Bacterial, but not fungal, communities show spatial heterogeneity in European beech (Fagus sylvatica L.) deadwood. FEMS Microbiology Ecology, 99(4), fiad023.

Bulygina, O.N., Razuvaev, V.N., Trofimenko, L.T., Shvets, N.V., 2022. Opisanie massiva dannyh srednemesyachnoy temperatury vozduha na stantsiyah Rossii. Svidetelstvo o gosudarstvennoy registratsii bazy dannyh No 2014621485. URL: http://meteo.ru/data/156-temperature (data access 25.10.2022) – Description of the data array of average monthly air temperature at stations in Russia. Certificate of state registration of the database No 2014621485.

Bunnell, F.L., Houde, I., Johnston, B., and Wind, E. 2002. How dead trees sustain live organisms in western forests. In Proceedings of the Symposium on the Ecology and Management of Dead Wood in Western Forests, 2–4 November 1999, Reno, Nevada. Technical coordinators: W.F. Laudenslayer, Jr., P.J. Shea, B.E. Valentine, C.P. Weatherspoon, and T.E. Lisle. USDA For. Serv. Gen. Tech. Rep. PSW-GTR-181. pp. 291–318.

Callahan B.J., McMurdie P.J., Rosen M.J., Han A.W., Johnson A.J.A., Holmes S.P. 2016. DADA2: High-resolution sample inference from Illumina amplicon data. Nat. Methods. 13, 581–583. 10.1038/nmeth.3869

Chao, A., Gotelli, N. J., Hsieh, T. C., Sander, E. L., Ma, K. H., Colwell, R. K., & Ellison, A. M. (2014). Rarefaction and extrapolation with Hill numbers: a framework for sampling and estimation in species diversity studies. Ecological monographs, 84(1), 45–67.

Chytrý, M., Tichý, L., Hennekens, S.M., et al., 2020. EUNIS Habitat Classification: expert system, characteristic species combinations and distribution maps of European habitats. Appl. Veg. Sci. 23, 3. 10.1111/avsc.12519.

Fernandez, C. W., Koide, R. T. (2013). The function of melanin in the ectomycorrhizal fungus *Cenococcum geophilum* under water stress. Fungal Ecology, 6(6), 479–486.

Fierer, N, Jackson, R.B. The diversity and biogeography of soil bacterial communities. Proceedings of the National Academy of Sciences. 2006. 103 (3) 626–631. 10.1073/pnas.0507535103

Guhr, A., Borken, W., Spohn, M., Matzner, E. Redistribution of soil water by a saprotrophic fungus enhances carbon mineralization. PNAS. 2015. 112(47): 14647–14651. 10.1073/pnas.1514435112

Girvan, M. S., Bullimore, J., Pretty, J. N., Osborn, A. M., & Ball, A. S. (2003). Soil type is the primary determinant of the composition of the total and active bacterial communities in arable soils. Applied and Environmental Microbiology, 69(3), 1800–1809.

Hemkemeyer M., Dohrmann A.B., Christensen B.T., Tebbe C.C. 2018. Bacterial preferences for specific soil particle size fractions revealed by community analyses. Front. Microbiol. 9, 149. 10.3389/fmicb.2018.00149

Hendershot, W. H., & Duquette, M. (1986). Exchangeable cation and effective CEC by the BaCl2 method. Soil Sampling and Methods of Analysis, 168–170.

Hsieh, T. C., Ma, K. H. F., & Chao, A. (2016). iNEXT: an R package for rarefaction and extrapolation of species diversity (H ill numbers). Methods in ecology and evolution, 7(12), 1451–1456.

Ilyin, B.M., Bulygina, O.N., Bogdanova, E.G., Veselov, V.M., Gavrilova, S.Yu., 2022. Opisanie massiva mesyachnyh summ osadkov, s ustraneniem sistematicheskih pogreshnostej osadkomernyh priborov. URL: http://meteo.ru/data/506-mesyachnye-summy-osadkov-s-ustraneniem-sistematicheskikh-pogreshnostej-osadkomernykh-priborov (data access 25.10.2022) Description of the array of monthly precipitation sums, with the elimination of systematic errors of precipitation gauges.

IUSS Working Group WRB. 2022. World Reference Base for Soil Resources. International Soil Classification System for Naming Soils and Creating Legends for Soil Maps, 4th ed. Vienna, Austria: International Union of Soil Sciences (IUSS).

Jany, J.L., Martin, F., Garbaye, J. (2003). Respiration activity of ectomycorrhizas from Cenococcum geophilum and Lactarius sp. in relation to soil water potential in five beech forests. Plant and Soil, 255(2), 487–494.

Jiao J.-Y., Liu L., Hua Z.-S., Fang B.-Z., Zhou E.-M., Salam N., Hedlund B.P., Li W.-J. 2021. Microbial dark matter coming to light: challenges and opportunities. NSR. 8 (3), nwaa280. 10.1093/nsr/nwaa280

Johnston S.R., Boddy L., Weightman A.J. 2016. Bacteria in decomposing wood and their interactions with wood-decay fungi. FEMS Microbiol. Ecol. 92 (11), fiw179. 10.1093/femsec/fiw179

Kersters K., De Vos P., Gillis M., Swings J., Vandamme P., Stackebrandt E. 2006. Introduction to the Proteobacteria. In The Prokaryotes. Dworkin M., Falkow S., Rosenberg E., Schleifer K.H., Stackebrandt E., Eds. New York: Springer, pp. 3–37. 10.1007/0-387-30745-1_1

Khanina L., Bobrovsky M. Changes in soil texture under deadwood in broad-leaved forests // Russian Journal of Ecosystem Ecology. 2024. Vol. 9 (4). DOI: 10.21685/2500-0578-2024-4-1

Khanina L., Bobrovsky M., Smirnov V., Romanov M. 2023. Wood decomposition, carbon, nitrogen, and pH values in logs of 8 tree species 14 and 15 years after a catastrophic windthrow in a mesic broad-leaved forest in the East European plain. For. Ecol. Manage. 545, 121275. 10.1016/j.foreco.2023.121275

Khanina L.G., Ivashchenko K.V., Smirnov V.E., Bobrovskii M.V. 2024. Carbon and nitrogen stocks and microbial activity of the humus horizon of loamy soils after mass windthrow in the broadleaved forest of the Kaluzhskie Zaseki Nature Reserve. Eurasian Soil Sci. 57 (11), 1806–1818. 10.1134/S1064229324601720

Kielak A.M., Scheublin T.R., Mendes L.W., Van Veen J.A., Kuramae E.E. 2016. Bacterial community succession in pine wood decomposition. Front. Microbiol. 7, 231. 10.3389/fmicb.2016.00231

Kuzyakov, Y., & Blagodatskaya, E. (2015). Microbial hotspots and hot moments in soil: Concept & review. Soil biology and biochemistry, 83, 184–199.

Lasota, J., Piaszczyk, W., & Błońska, E. (2022). Fine woody debris as a biogen reservoir in forest ecosystems. Acta Oecologica, 115, 103822.

Lindenmayer D.B., Franklin J.F. 2002. Conserving Forest Biodiversity: A Comprehensive Multiscaled Approach. London: Island Press.

Lladó S., López-Mondéjar R., Baldrian P. 2017. Forest soil bacteria: diversity, involvement in ecosystem processes, and response to global change. Microbiol. Mol. Biol. Rev. 81 (2), e00063–16. 10.1128/mmbr.00063-16

Lorch, J. M., Lindner, D. L., Gargas, A., Muller, L. K., Minnis, A. M., & Blehert, D. S. (2013). A culture-based survey of fungi in soil from bat hibernacula in the eastern United States and its implications for detection of Geomyces destructans, the causal agent of bat white-nose syndrome. Mycologia, 105(2), 237–252.

Lukina N.V., Bartalev S.A., Geraskina A.P., Plotnikova A.S., Gornov A.V., Ershov D.V., Gavrilyuk E.A., Kuznetsova A.I., Shevchenko N.E., Tikhonova E.V., Danilova M.A., Tebenkova D.N., Smirnov V.E., Ruchinskaya E.V. 2023. Role of old-growth forests in carbon accumulation and storage. *Izv. Ross. Akad. Nauk*, Ser. Geogr. 87 (4), 536–557. 10.31857/S2587556623040064

von Lützow, M., Kögel-Knabner, I., Ekschmitt, K., Matzner, E., Guggenberger, G., Marschner, B., & Flessa, H. (2006). Stabilization of organic matter in temperate soils: mechanisms and their relevance under different soil conditions - a review. European Journal of Soil Science, 57(4), 426–445.

Mäkipää, R., Rajala, T., Schigel, D., Rinne, K. T., Pennanen, T., Abrego, N., & Ovaskainen, O. (2017). Interactions between soil-and dead wood-inhabiting fungal communities during the decay of Norway spruce logs. The ISME journal, 11(9), 1964–1974.

McDonald D., Jiang Y., Balaban M. et al. 2024. Greengenes2 unifies microbial data in a single reference tree. Nat. Biotechnol. 42, 715–718. 10.1038/s41587-023-01845-1

McMurdie P.J., Holmes S. 2013. An R package for reproducible interactive analysis and graphics of microbiome census data. PLoS One. 8 (4), e61217. 10.1371/journal.pone.0061217.

Meena M., Yadav G., Sonigra P., Nagda A., Mehta T., Swapnil P., Harish, Marwal A., Kumar S. 2023. Multifarious responses of forest soil microbial community toward climate change. Microb. Ecol. 86 (1), 49–74. 10.1007/s00248-022-02051-3

Minnich C., Peršoh D., Poll C., Borken W. 2021. Changes in chemical and microbial soil parameters following 8 years of deadwood decay: an experiment with logs of 13 tree species in 30 forests. Ecosystems. 24, 955–967. 10.1007/s10021-020-00562-z

Moll, J., Kellner, H., Leonhardt, S., Stengel, E., Dahl, A., Bässler, C., … & Hoppe, B. (2018). Bacteria inhabiting deadwood of 13 tree species are heterogeneously distributed between sapwood and heartwood. Environmental Microbiology, 20(10), 3744–3756.

Muhr J., Messier C., Delagrange S., Trumbore S., Xu X., Hartmann H. 2016. How fresh is maple syrup? Sugar maple trees mobilize carbon stored several years previously during early springtime sap-ascent. New Phytol. 209 (4), 1410–1416. 10.1111/nph.13782

Nannipieri, P., Giagnoni, L., Landi, L., & Renella, G. (2012). Role of phosphatase enzymes in soil. In Phosphorus in Action (pp. 215–243). Springer, Berlin, Heidelberg.

Novenko E.Yu., Tsyganov A.N., Mazei N.G., Kupriyanov D.A., Rudenko O.V., Bobrovsky M.V., Erman N.F., Nizovtsev V.A. Palaeoecological evidence for climatic and human impacts on vegetation in the temperate deciduous forest zone of European Russia during the last 4200 years: A case study from the Kaluzhskiye Zaseki Nature Reserve // Quaternary International. 2019. Vol. 516. P. 58-69. DOI: 10.1016/j.quaint.2018.06.028

Oksanen J., Simpson G., Blanchet F., Friendly M., Kindt R., Legendre P., McGlinn D., Minchin P.R., O’Hara R.B., Simpson G.L., Solymos P., Stevens M.H.H., Szoecs E., Wagner H. 2025. Vegan: Community Ecology Package. 10.32614/CRAN.package.vegan

Pavlopoulos G.A., Baltoumas F.A., Liu S., Selvitopi O., Camargo A.P., Nayfach S., Azad A., Roux S., Call L., Ivanova N.N., Min Chen I., Paez-Espino D., Karatzas E., Novel Metagenome Protein Families Consortium, Iliopoulos I., Konstantinidis K., Tiedje J.M., Pett-Ridge J., Baker D., Visel A., Ouzounis C.A., Ovchinnikov S., Buluç A., Kyrpides N.C. 2023. Unraveling the functional dark matter through global metagenomics. Nature. 622 (7983), 594–602. 10.1038/s41586-023-06583-7

Peter, M., Kohler, A., Ohm, R. A., Kuo, A., Krützmann, J., Morin, E., … & Martin, F. M. (2016). Ectomycorrhizal ecology is imprinted in the genome of the dominant symbiotic fungus *Cenococcum geophilum*. Nature communications, 7(1), 12662.

Pioli, S., Clagnan, E., Chowdhury, A. A., Bani, A., Borruso, L., Ventura, M., … & Brusetti, L. (2023). Structural and functional microbial diversity in deadwood respond to decomposition dynamics. Environmental Microbiology, 25(11), 2351–2367.

Põlme, S., Abarenkov, K., Henrik Nilsson, R., Lindahl, B. D., Clemmensen, K. E., Kauserud, H., … & Tedersoo, L. (2020). FungalTraits: a user-friendly traits database of fungi and fungus-like stramenopiles. Fungal diversity, 105(1), 1–16.

Popadiouk, R.V., Smirnova, O.V., Zaugolnova, L.B., Khanina, L.G., Bobrovsky, M.V., Yanitskaya, T.Y., 1999. Zapovednik “Kaluzhskie Zaseki”, in: Smirnova, O.V., Shaposhnikov, E.S. (Eds.), Suktsessionnye Processy v Zapovednikakh Rossii i Problemy Sokhraneniya Biologicheskogo Raznoobraziya. Russkoe Botanicheskoe Obshchestvo, SPb., s. 58–105 – The Kaluzhskie zaseki State Nature Reserve (in Russian).

Qi L., Yuan J., Zhang W., Liu H., Li Z., Bol R., Zhang S. 2023. Metagenomics reveals the underestimated role of bacteria in the decomposition of downed logs in forest ecosystems. Soil Biol. Biochem. 187, 109185. 10.1016/j.soilbio.2023.109185

R Core Team 2025. R: A Language and Environment for Statistical Computing. Vienna: R Foundation for Statistical Computing. https://www.R-project.org.

Rakitin A.L., Kulichevskaya I.S., Beletsky A.V., Mardanov A.V., Dedysh S.N., Ravin N.V. 2024. Verrucomicrobia of the family *Chthoniobacteraceae* participate in xylan degradation in boreal peat soils. Microorganisms. 12 (11), 2271. 10.3390/microorganisms12112271

Rice, A. V., & Currah, R. S. (2006). Two new species of *Pseudogymnoascus* with *Geomyces* anamorphs and their phylogenetic relationship with *Gymnostellatospora*. Mycologia, 98(2), 307–318.

Rousk, J., Bååth, E., Brookes, P. C., et al. (2010). Soil bacterial and fungal communities across a pH gradient. Soil Biology and Biochemistry, 42(6), 920–929.

Savelyev D. 2025. Data on composition of soil bacteria under deadwood of various tree species after windthrow in old-growth broad-leaved forests: figshare. Dataset. https://figshare.com/articles/dataset/_/30593849

Savelyev D.S., Shelyakin P.V., Tutukina M.N., Bessonova T.A., Bobrovsky M.V., Khanina L.G. Changes in Bacterial Diversity and Bacterial DNA Load in The Soil Beneath Deadwood of Various Tree Species Following Windthrow in Old-Growth Broadleaved Forests // Molecular Biology. 2026. Vol. 60. No. 4. P. 589–603.

Schimel D.S., Braswell B.H., Holland E.A., McKeown R., Ojima D.S., Painter T.H., Parton W.J., Townsend A.R. 1994. Climatic, edaphic, and biotic controls over storage and turnover of carbon in soils. Global Biogeochem. Cycles. 8, 279–293. 10.1029/94GB00993

Schmidt, M. W., Torn, M. S., Abiven, S., Dittmar, T., Guggenberger, G., Janssens, I. A., … & Trumbore, S. E. (2011). Persistence of soil organic matter as an ecosystem property. Nature, 478(7367), 49–56.

Seaton F.M., George P.B., Lebron I., Jones D.L., Creer S., Robinson D.A. 2020. Soil textural heterogeneity impacts bacterial but not fungal diversity. Soil Biol. Biochem. 144, 107766. 10.1016/j.soilbio.2020.107766

Shashkov, M.P., Bobrovsky, M.V., Shanin, V.N., et al., Data on 30-year stand dynamics in an old-growth broad-leaved forest in the Kaluzhskie Zaseki State Na ture Reserve, Russia, Nat. Conserv. Res., 2022, vol. 7, Suppl. 1, pp. 24–37.

Smirnov V.E., Khanina L.G., Ivashchenko K.V., Bobrovsky M.V. 2024. Assessment of the impacts of large downed oak trunks on soil properties based on mixed-effects statistical models. In Proc. Int. Conf. “Mathematical Biology and Bioinformatics.” Lakhno V.D., Ed. Pushchino: IMPB RAS. vol. 10. Paper no. e52. 10.17537/icmbb24.54

Smirnova O.V., Bobrovsky M.V., Khanina L.G., Braslavskaya T.Yu., Starodubtseva E.A., Evstigneev O.I., Korotkov V.N., Smirnov V.E., Ivanova N.V. 2017. Nemoral forests. In European Russian Forests. Their Current State and Features of Their History. Smirnova O.V., Bobrovsky M.V., Khanina L.G., Eds. Netherlands: Springer, pp. 337–480. 10.1007/978-94-024-1172-5

Sollins, P., Homann, P., & Caldwell, B. A. (1996). Stabilization and destabilization of soil organic matter: mechanisms and controls. Geoderma, 74(1-2), 65–105.

Solomeshch, A. I., Grigoriev, I. N., Khaziakhmetov, R. M., & Baisheva, E. Z. (1993). Sintaksonomiya lesov Yuzhnogo Urala. V. Khvoyno-shirokolistvennye lesa [Syntaxonomy of forests of the Southern Urals. V. Coniferous-broadleaved forests] (Manuscript Deposited at VINITI 02.08.1993, No. 1464-V93). Ufa. [in Russian]

Steinebrunner, F., Tischer, A., Medicus, T., Huth, F., & Bernhardt-Römermann, M. (2025). The effects of deadwood on tree regeneration and microsites: A systematic review. Forest Ecology and Management, 596, 123096.

Stokland J.N., Siitonen J., Jonsson B.G. 2012. Biodiversity in Dead Wood. Cambridge: Cambridge Univ. Press. 10.1017/CBO9781139025843

Stutz, K. P., Dann, D., Wambsganss, J., Scherer-Lorenzen, M., & Lang, F. (2017). Phenolic matter from deadwood can impact forest soil properties. Geoderma, 288, 204–212.

Tláskal, V., & Baldrian, P. (2021). Deadwood-inhabiting bacteria show adaptations to changing carbon and nitrogen availability during decomposition. Frontiers in Microbiology, 12, 685303.

Tláskal, V., Zrůstová, P., Vrška, T., & Baldrian, P. (2017). Bacteria associated with decomposing dead wood in a natural temperate forest. FEMS Microbiology Ecology, 93(12), fix157.

Uroz S., Ioannidis P., Lengelle J., Cebron A., Morin E., Buéé M., Martin F. 2013. Functional assays and metagenomic analyses reveals differences between the microbial communities inhabiting the soil horizons of a Norway spruce plantation. PLoS One. 8 (2), e55929. 10.1371/journal.pone.0055929

de Vries F.T., Manning P., Tallowin J.R., Mortimer S.R., Pilgrim E.S., Harrison K.A., Tallowin J.R.B., Mortimer S.R., Pilgrim E.S., Harrison K.A., Hobbs P.J., Quirk H., Shipley B., Cornelissen J.H.C., Kattge J., Bardgett R.D. 2012. Abiotic drivers and plant traits explain landscape-scale patterns in soil microbial communities. Ecol. Lett. 15 (11), 1230–1239. 10.1111/j.1461-0248.2012.01844.x

Wang Q., Wang Z., Penuelas J., Cao R., Wang L., Yang W. 2024. Decaying logs and gap positions jointly mediate the structure and function of soil bacterial community in the forest ecosystem. For. Ecol. Manage. 567, 122070. 10.1016/j.foreco.2024.122070

Wang Z., Xu M., Li F., Bai Y., Hou J., Li X., Cao R., Deng Y., Jiang Y., Wang H., Yang W. 2023. Changes in soil bacterial communities and functional groups beneath coarse woody debris across a subalpine forest successional series. Global Ecol. Conserv. 43, e02436. 10.1016/j.gecco.2023.e02436

Zaugol’nova, L.B. and Braslavskaya, T.Yu., The analysis of mesophytic broad-leaved forest associations in the center of European Russia, Rastit. Ross., 2003, no. 4, pp. 3–28.

Zhu, M., You, W., & Ni, S. (2026). Deadwood-driven improvements in soil health are amplified by forest gaps and elevation. CATENA, 272, 110349.

Žifčáková, L., Větrovský, T., Howe, A., & Baldrian, P. (2016). Microbial activity in forest soil reflects the changes in ecosystem properties between summer and winter. Environmental microbiology, 18(1), 288–301.

